# Control over a mixture of policies determines change of mind topology during continuous choice

**DOI:** 10.1101/2024.04.18.590154

**Authors:** Justin M. Fine, Seng-Bum Michael Yoo, Benjamin Y. Hayden

## Abstract

Behavior is naturally organized into categorically distinct states with corresponding patterns of neural activity; how does the brain control those states? We propose that states are regulated by specific neural processes that implement meta-control that can blend simpler control processes. To test this hypothesis, we recorded from neurons in the dorsal anterior cingulate cortex (dACC) and dorsal premotor cortex (PMd) while macaques performed a continuous pursuit task with two moving prey that followed evasive strategies. We used a novel control theoretic approach to infer subjects’ moment-to-moment latent control variables, which in turn dictated their blend of distinct identifiable control processes. We identified low-dimensional subspaces in neuronal responses that reflected the current strategy, the value of the pursued target, and the relative value of the two targets. The top two principal components of activity tracked changes of mind in abstract and change-type-specific formats, respectively. These results indicate that control of behavioral state reflects the interaction of brain processes found in dorsal prefrontal regions that implement a mixture over low-level control policies.

## INTRODUCTION

As humans and other animals interact with the world, we move between discrete behavioral states that influence how we respond to stimuli in our environment (Anderson and Perona, 2014; Brown and De Bivort, 2014; Periera et al., 2020). Neuroscientists have begun to identify the neural correlates of these states and to delineate their effects on neural activity (Marques et al., 2020; Calhoun et al., 2019; Markowitz et al., 2018; Voloh et al., 2023). These findings raise the question of how the brain selects between possible behavioral states on a moment-to-moment basis. Resource-limited (including time-limited) agents must solve a meta-control problem of allocating control between different goals or tasks, which can involve blended strategies (Lieder and Griffiths, 2017; Yeung and Summerfield, 2012). A classic expression of such state control is a change-of mind in choice or task-switching (Fleming et al., 2018; Kiani et al., 2014; Sarafyazd & Jazayeri, 2019). However, unlike static choice tasks, in dynamic interactive environments, meta-control requires integrating multimodal information continuously over time and dynamically weighing the costs and benefits of switching while continuing to act. In other words, to choose between strategies, we must combine and deploy multiple action control-policies on the fly.

## RESULTS

To gain insight into how the brain controls behavioral states in dynamic contexts, we developed a prey-pursuit task in which macaque subjects moved a joystick to pursue one of two moving targets (Yoo et al., 2020 and 2021, **Figure 1A**). This task involved two conditions, a one-prey and a two-prey condition; due to our focus here on strategic switching, in the present study we focus on the two-prey condition. Prey differed on the dimension of reward value and speed (**Figure 1B**). Subjects performed quite well in this task; they successfully captured the prey in ∼80% of trials (subjects H: 84.9%; subject K: 79.0%). The average time for capturing a prey item was 3.87 s (H: 3.21; K: 4.53). Capture time was roughly the same for all reward differences between prey items, although subject H was slightly faster in the smallest difference condition (3.19 s; *F* = 3.04, *p* = 0.028).

**Figure 1.**
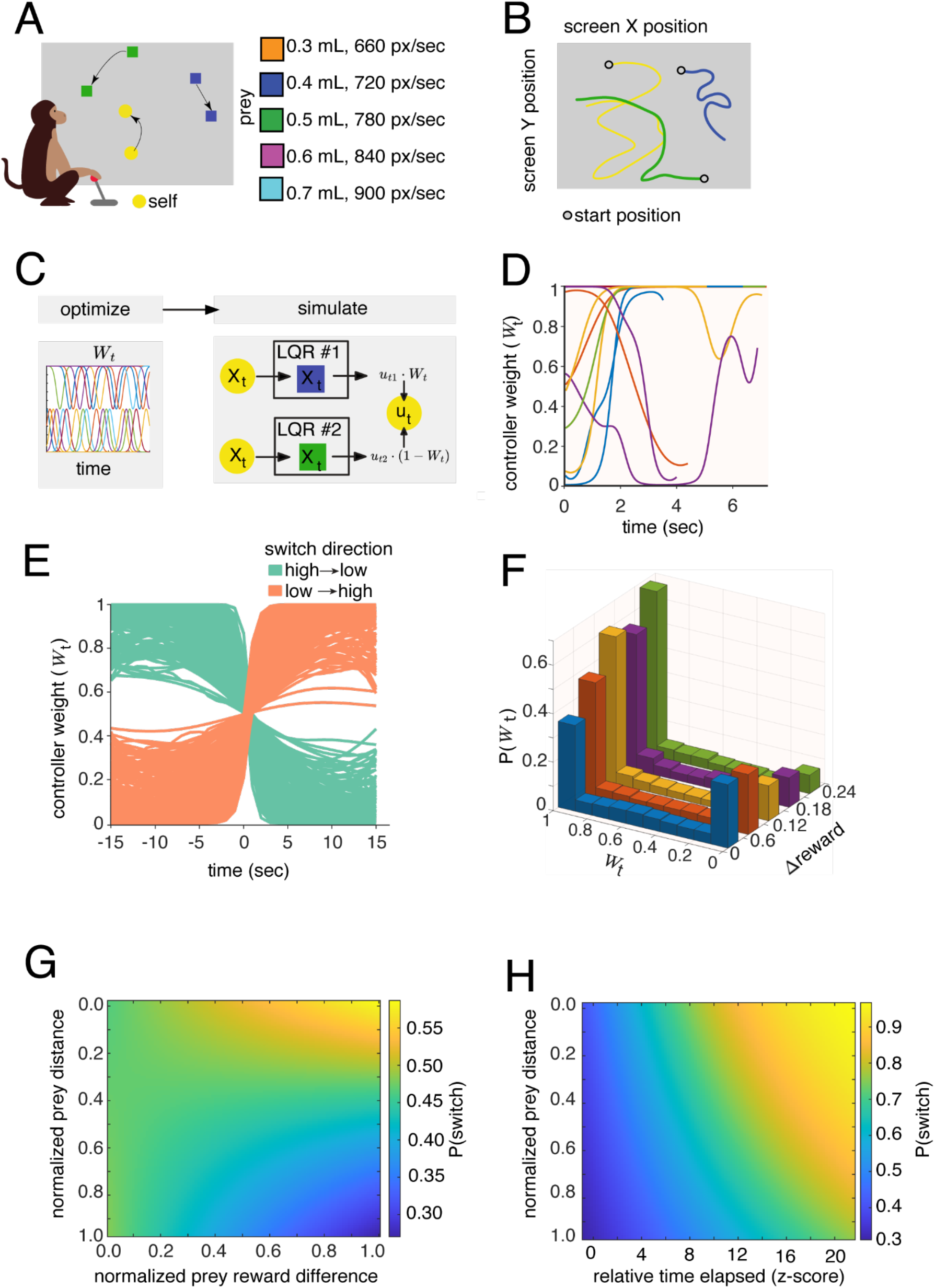
**A**. Illustration of prey-pursuit task. A macaque subject looks at a computer monitor that portrays shapes representing the self avatar (circle), and one or two prey (squares). The different square colors indicate different possible prey values (juice volume). **B**. Example movement traces of subject chasing two possible prey in a trial, with the circle representing the start point. **C**. Graphical depiction of our control-theory based decomposition of pursuit actions. For each trial, and optimization iteration, we model the latent pursuit (i.e., LQR policy mixture) variable W_*t*_ with radial basis functions over the trial time, alongside optimizing LQR parameters. These radial bases weights are combined to create a singular W_*t*_ time series, which is then used as a proposed mixture of LQR control signals (per prey target). The predicted control *u* is then compared to the subject’s actual velocity, and error is computed and cycled through optimization (full details in methods). **D**. Example controller W_*t*_ found for several trials (different color traces) for subject H. **E**. For finding the switch points, we searched trials for a crossing of W_*t*_ = 0.5. When time-locking the W_*t*_ time series to the W_*t*_ = 0.5, we can find stereotyped sigmoidal W_*t*_ trajectories depicting whether the subject (here, H) made switches from a high to low (green) value target or low to high (orange). These traces represent all the switches from one session from subject H. **F**. Distributions of whole trial W_*t*_ for both subjects, as a function of the prey reward difference (Δreward). As Δreward increased, we see an expected shift from mostly bimodal pursuit of targets (Δreward =0.6) to a near unimodal focus of subjects’ pursuit on the high reward target when Δreward was high (0.24). **G**. Predicted probability of a switch, P(switch), landscape as a function of the interaction between distance of the switched-to-prey and Δreward. **H**. Predicted probability of a switch, P(switch), landscape as a function of the interaction between distance of the switched-to-prey and relative trial time elapsed (both subjects). Longer trials predicted higher P(switch).

To identify the pursued target at every point in time, we inferred a latent continuous weighting variable (W_*t*_) that indicates which of two prey the subject was pursuing at any given time. Inferring this variable is non-trivial. A strategy that uses linear algebraic decomposition of kinematics (e.g., scalar projections) would necessarily assume that the weighting of targets is isomorphic, that is, that greater velocity towards target 1 (relative to target 2) implies pursuit is towards target 1. This relationship is not generally true, even in the simple case of two targets (see **Methods**). Instead, the meta-control signal, the drivers of the agent’s instantaneous control, is a latent variable that must be inferred. At the same time, approaches like Hidden Markov Modeling (HMM) assume choice weighing is a categorical variable rather than continuous, and do not reflect an underlying model. For these reasons, HMM-like approaches are poorly matched for continuous domains and are particularly ill-suited to uncovering meta-control signals that encode continuous mixtures over low-level control policies. Consequently, these approaches often fail to detect clear changes of mind (**Supplementary Result S1**). To uncover this continuous mixing control signal, we developed a novel model-based decomposition that leverages control theory. Our approach follows from an optimization principle that control in a nonlinear cost landscape can be approximated by a compositional mixture of linear, optimal controllers (Todorov, 2009; Dvijotham and Todorov, 2012).

For each trial, we combine two infinite-horizon, linear-quadratic regulator (LQR) controllers and optimize a normalized instantaneous weighting over the controllers (W_*t*_, **Figure 1C**, see **Methods** for details), alongside the controller gains. The weight on each controller dictates how much a subject’s instantaneous control (i.e., velocity) is attributable to each prey’s specific LQR (**Figure 1D**). The latent controller weight trajectory is estimated with basis functions (of fixed but optimized width) that tessellate trial time. This approach performs substantially better than algebraic projection and HMM methods in ground-truth model recovery (**Supplementary S1.2-S1.3**). To validate our approach, we generated simulated data using ground-truth W_*t*_ functions (**Table S1**) to control two LQR controllers and used our method to recover W_*t*_ values. Confirming the validity of this approach, we recovered simulated W_*t*_ signals (the inferred W_*t*_ variable) with high fidelity, regardless of complexity (*r* = 0.92 ± 0.08); algebraic projection (*r* = 0.32 ± 0.18) and HMMs (*r* = 0.34 ± 0.04) fared similarly to one another, with both being significantly worse than our model (**Table S1, Figure S2-S3**).

We can use the recovered W_*t*_ trajectories to understand subjects’ behavior (**Figure 1E-F**). We defined pursuit of the larger reward as moments when W_*t*_>0.85 and pursuit of the lower reward as moments when W_*t*_<0.15. Both subjects spent an average of ∼55% of their time pursuing the higher-value reward and ∼15% pursuing the lower-value target (t-tests comparing time for higher-versus lower-value prey; H: t = 23.92, *p* < 0.0001, K: t =21.34, *p* < 0.0001). The rest of the time (∼30%) was spent with W_*t*_ parameters between these two extrema (i.e., 0.15<W_*t*_<0.85)).

Both subjects tended to focus on a single target at each timepoint, rather than a strategically pursue both prey (cf. Cisek, 2012). W_*t*_ distributions showed that subjects become increasingly likely to pursue a higher-value reward target as differences in prey value increase (illustrative distributions for subject H, **Figure1E**), consistent with a softmax strategy across reward prey differences. Subjects presumably favor a more unimodal strategy with increasing reward difference to maximize their chances of obtaining a high-value reward.

Our identification of inferred weight values allows analysis of switch points between pursuit states (**Figure 1F**). Both subjects made a moderate number of switches (subject H: 28% of trials; subject K: 51% of trials). We identified the factors that predicted switches using a Lasso logistic regression (see **Methods** and **Table S2** for all predictors). We found several significant predictors (**Figure S4**). Some of these included trial time elapsed (log-odds CI: [0.13, 0.17]), and increased prey reward differences (log-odds CI: [-0.20, -0.06]). In terms of prey distances leading up to a switch, we found the closer, on the computer monitor, the switched-to-prey was to the subject’s avatar, the more likely a switch (log-odds CI: [0.04, 0.15]). These core factors all interacted, to reveal a complex pattern of switching behavior. For example, distance to switched prey and time elapsed (log-odds CI: [-0.04, -0.009], Figure 1G) exerted a nonlinear influence on switching, as well as distance to switched prey and delta reward (log-odds CI: [-0.36, -0.21, Figure 1H]) interacted to determine the switch behavior. This indicates subjects were agnostic to the switched prey distance when the difference in reward was zero (Figure 1G) or little time had elapsed (Figure 1H). Strategically, the effective value prey proximity had on switching was dependent on internal (value difference) and external factors (time elapsed).

To elucidate how these policy control decisions are implemented neurally, we examined responses of neurons collected from the dorsal anterior cingulate cortex (dACC) in both subjects (H: n=119, K: n=48, **Figure 2A**). Our previous work indicated that these neurons are selective to the position and speed of the subject’s avatar, among other variables (Yoo et al, 2020 and 2021). Here, we asked if dACC neurons encode the latent decision weight (W_*t*_, **Figure 2B**). We examined single-neuron variable encoding using a linear regression (false discovery rate *=* 1x10^-5^) for each neuron. In both subjects, we find a high proportion of neurons were tuned to W_*t*_ (all W_*t*_ combinations: H: 49% and K: 71%). We show W_*t*_ tuning curves for two illustrative neurons in **Figure 2C and D**. The proportion of neurons whose responses were selective to W_*t*_ was essentially the same as the proportion tuned to speed or reward difference (**Figure 2B**). Moreover, W_*t*_ tuning was both linearly and nonlinearly mixed with coding of prey value difference and self-speed, indicating a general tuning distribution of mixed selectivity (**Figure 2B**, Rigotti et al., 2013).

**Figure 2.**
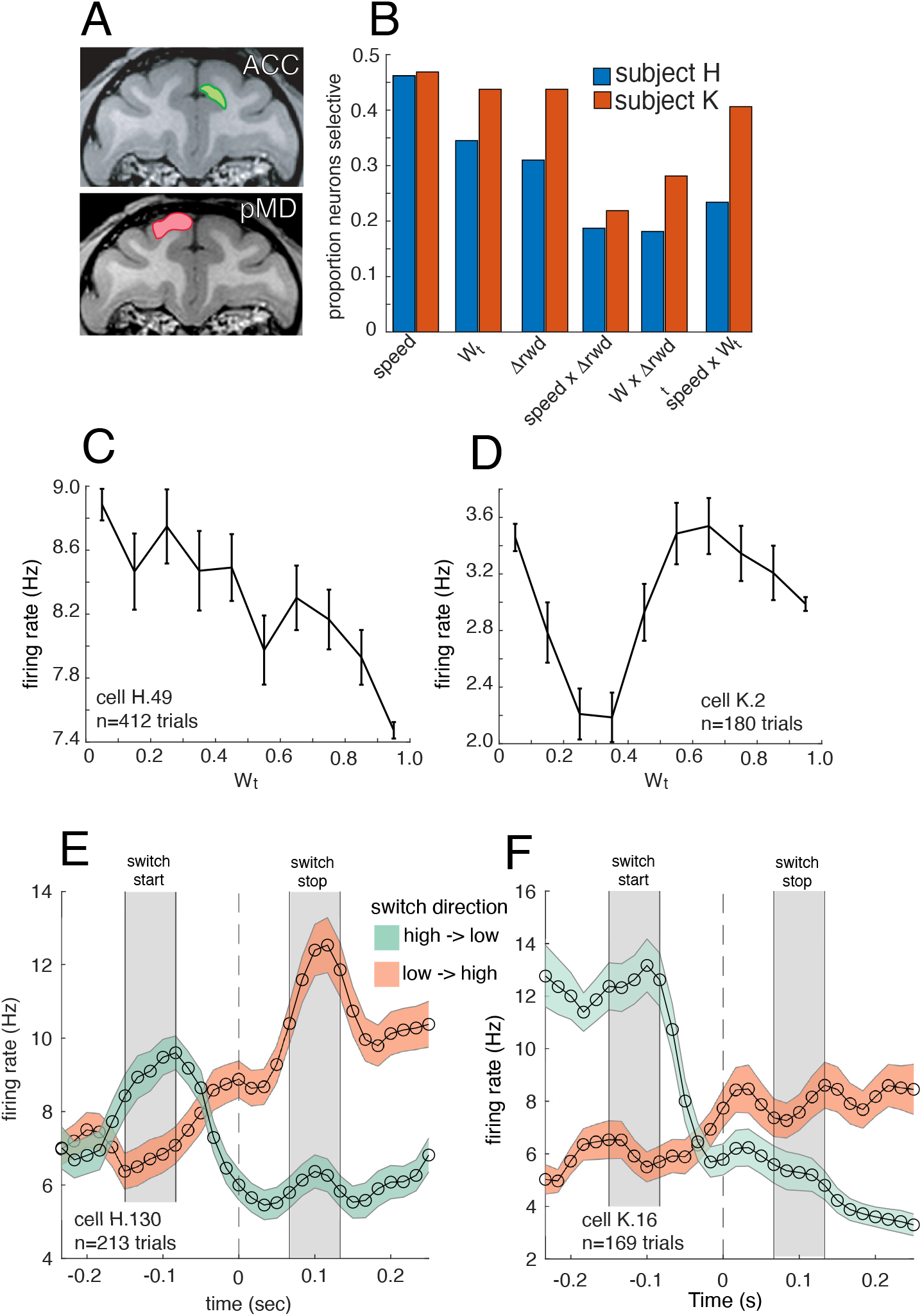
**A**. MRI sections showing average recording for both subjects in ACC (top) and subject H’s PMd. **B**. Results of selectivity analysis for both subjects’ ACC neurons, showing their proportion of neurons selective to relevant task-variables across a whole experimental session. **C-D**. Tuning curves for two ACC neurons with significant Wt tuning. **E-F**. Firing rate profiles (mean +-standard error) for two neurons time-locked to the switch window (time=0; W_*t*_ =0.5). The neuron profiles are shown for switches that go from a high Wt (∼0.9) to a low W_*t*_ (∼0.1) in green, and low to high switches are shown in pink. Because the data are time-normalized or warped to the switch center (W_*t*_=0.5), the gray regions denote the median switch-start and stop-time ± 1 std.

What is distinct about online and continuous choice is that both acting and deciding effectively occur in parallel: subjects continue to chase the target while contemplating and then executing an impending switch (Yoo et al., 2020). Thus, we next asked how neural selectivity patterns support these choice switching processes. Using model-based derived switch points, we investigated how neuronal activity covaries with switches. Examining example neurons time-locked to the switch window, we see a pattern of heterogeneous responses to W_*t*_ (**Figure 2E-F**). Within these heterogeneous neuron profiles, is there a population code for W_*t*_? If neural populations indeed use a W_*t*_ representation, they should contain a readout that recapitulates the expected W_*t*_ behavioral trajectory (sigmoidal shape) during a switch from one target to another. Moreover, these neural W_*t*_ codes should be mirrored when comparing a switch from a high-to-low W_*t*_ versus a low-to-high W_*t*_. To quantify this, we used demixed PCA (dPCA) to identify separable projections of ACC population activity at the time of pursuit switches (Kobak et al., 2016). Specifically, we considered neural behavior around the time of switch points (that is, times when W_*t*_ crossed over the line of 0.5). We analyzed two types of switches to assess whether populations have a representation of W_*t*_: from the low to the high-value target, and vice versa (see **Methods**). Specifically, we aim to test a key prediction of the idea that subjects continuously mix policies. If a W_*t*_-like code is used, this predicts that a change-of-mind should have a low-dimensional code that approximates a sigmoidal trajectory from low to high W_*t*_ (and vice versa). We test this by comparing these two switch directions that start and end at different W_*t*_ values, we can ask whether there is a linear, demixable representation of W_*t*_.

The dPCA indicated 8 (out of 20) components accounted for >80% of variance. To examine linear decodability of W_*t*_ in the dACC population response, we projected held-out test trials onto the first two dPCA components (see **Methods**). Both components exhibited significant decoding of switch direction (that is, low-to-high vs high-to-low). In the ***first component*** (**Figure 3A** and **Figure S5A**), decoding of switch direction was possible across the whole switch window. This consistent decodability indicates information regarding switch direction information was linearly separable across the change-of-mind time-window. This component, then, could reflect a code that denotes a signal that could be used to separate differential causes for switching away from a high versus low value target.

**Figure 3.**
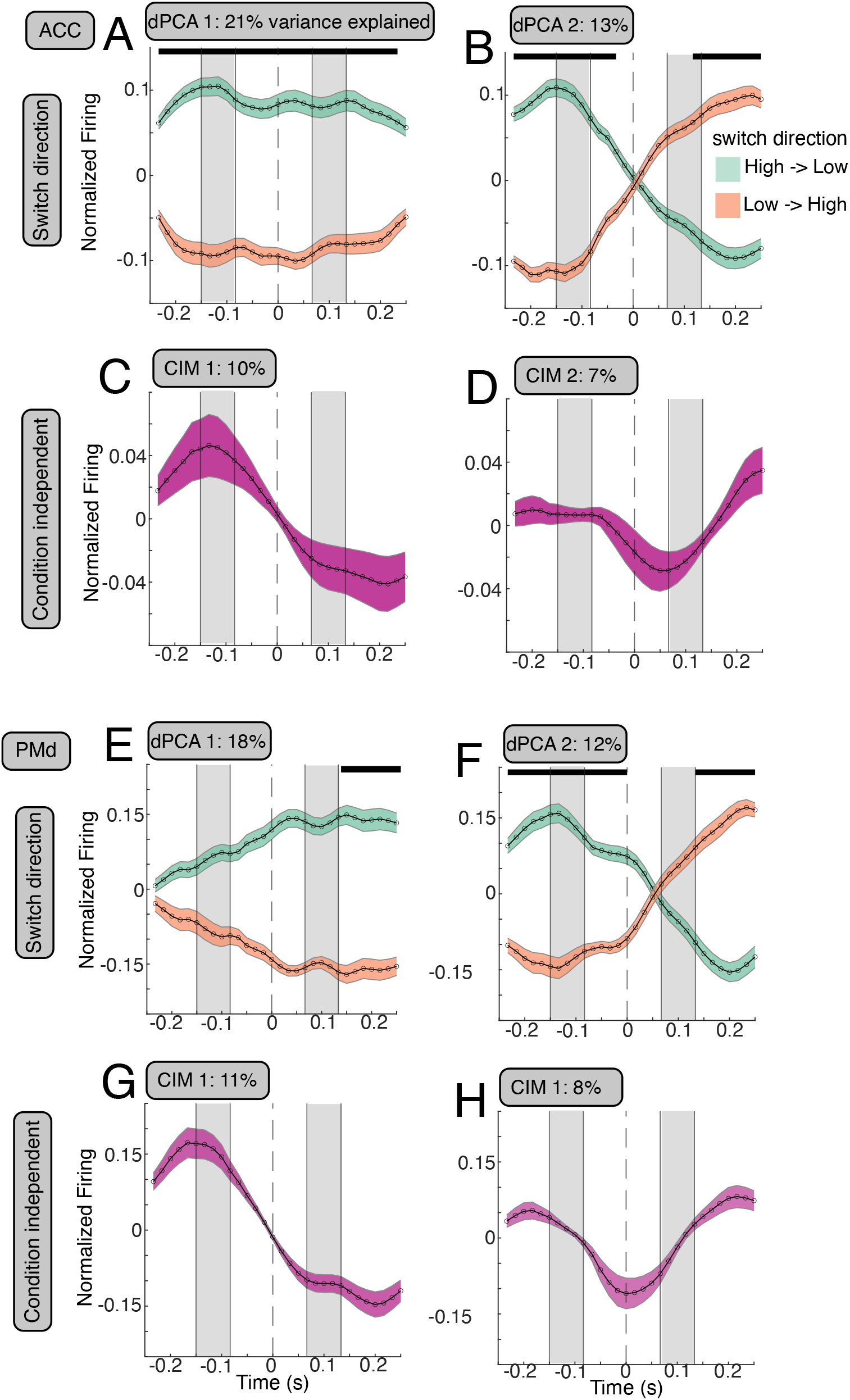
Recovered dPCA components presented as mean and standard error over their learned projections. Above each component, we show its ordered component number and % variance within that dPCA marginalization (either switch-direction or condition-independent). Black bars above the switch direction components indicate time-windows of significant decoding. Solid gray bars indicate the windows of switch -onset and -offset, in the form of median plus/minus standard deviation. The dashed bar indicates the Wt=0.5 switch point. **A-B**. First two switch direction components for ACC. **C-D**. First two CIM or condition-independent modes for ACC. **E-F**. First two switch direction components for PMd. **G-H**. First two CIM or condition-independent modes for PMd. The gray regions denote the median switch-start and stop-time ± 1 std..

The ***second component*** (**Figure 3B** and **Figure S5B**) has a sigmoidal switching shape consistent with W_*t*_ encoding. Specifically, this population projection shows a switch from low to high and high to low, while crossing at the time of the estimated switch point (W_*t*_ = 0.5). The dynamics of this component are consistent with implementing a linear code for W_*t*_ that may be used for control over control policy mixing and downstream motoric specification.

For comparison, we next examined neurons from dorsal premotor cortex (PMd, n=330 neurons total). These data were collected simultaneously with the dACC data in one subject. PMd selectivity (**Figure S6**) and dPCA (**Figure 3E-F**) response patterns generally matched those of ACC. Like dACC, in the first component we can see a code that separates the switch directions (see black decoding bars in **Figure 3E** and **Figure S5C**). Likewise, the PMd ***second component*** (**Figure 3F** and **Figure S5D**) was similar to the second component in dACC, mirroring a target pursuit weight code. Thus, it seems that both regions encode W_*t*_ in largely the same way.

The condition independent modes (CIM, equivalent to regression intercept terms), which are the dPCA modes independent of Wt, derived from the ACC and PMd dPCAs, coincided with key events during the switch. In both areas, this 1st component begins to change at the onset of switch, ramp during the switch window, and level-off at the end of switch (**Figure 3C, D, E, and H**). This pattern indicates a population code tracking the bookend switch events. The second CIMs exhibited a peak modulation at W_*t*_=0.5, indicating they could also mark change-of-mind progression, back into pursuit.

The idea that the brain may directly use a representation of W_*t*_ implies that the act of choosing which prey to pursue reflects dynamics that are shared across switch directions. One of our core hypotheses, grounded in the idea that W_*t*_ is a meta-control signal, is that a change of mind reflects a traversal of W_*t*_ by neural activity. The implication is that this activity evolves with some dynamics along a topology in a low-dimensional neural state space. One hypothesis this predicts is that traversal of W_*t*_ from either a low to high or high to low switch should use similar dynamics and that both conditions will reflect shared underlying dynamical systems.

An issue with comparing two putative dynamics systems to determine alignment is that state space geometry is not equivalent to topological (dynamics) conjugacy. Namely, two dynamic systems can have identical (or distinct) geometries yet have different (or similar) dynamics driving them (Galgali et al., 2023). For example, assuming both switch directions for W_*t*_ are reducible to linear systems, they could have very different eigenvalues that determine the dynamics, despite both being stable systems with similar geometries. A simple example in two-choice decision making is a 2D system driven to either choice by a line attractor, saddle point, or bistable discrete attactors will exhibit the same geometry on average (Ostrow et al., 2024).

We next asked whether the underlying dynamics between a change-of mind in either direction are similar. Namely, can two estimated systems underlying putative W*t* traversal be aligned in terms of their state spaces (i.e., vector fields; **Figure 4A**)? To quantify dynamical similarity, we used a recently derived metric (DSA; Ostrow et al., 2024; **Methods**). To estimate the dynamics, we used a dynamic mode decomposition (DMD) with unsupervised control estimation (Fieseler et al, 2020); the DMD method operates similarly to switching linear dynamical systems, by attempting to find unsupervised signals denoting changes in dynamics (Linderman et al., 2017). Separate DMDs were estimated for the same two switch directions used in dPCA; we used subsampled pseudo-populations to account for across-trial variation beyond the average dynamics (**Methods**; Galgali et al., 2023). First, we examined average subsampled dynamic mode decomposition for activity during the switch (**Methods**), for both switch directions (**Figure 4 D-E**). Both switch directions and brain areas’ average dynamics could be accounted for by a 5-dimensional system (>80% variance), with both areas exhibiting orthogonality in their geometry (**Figure 4 D-E**).

**Figure 4.**
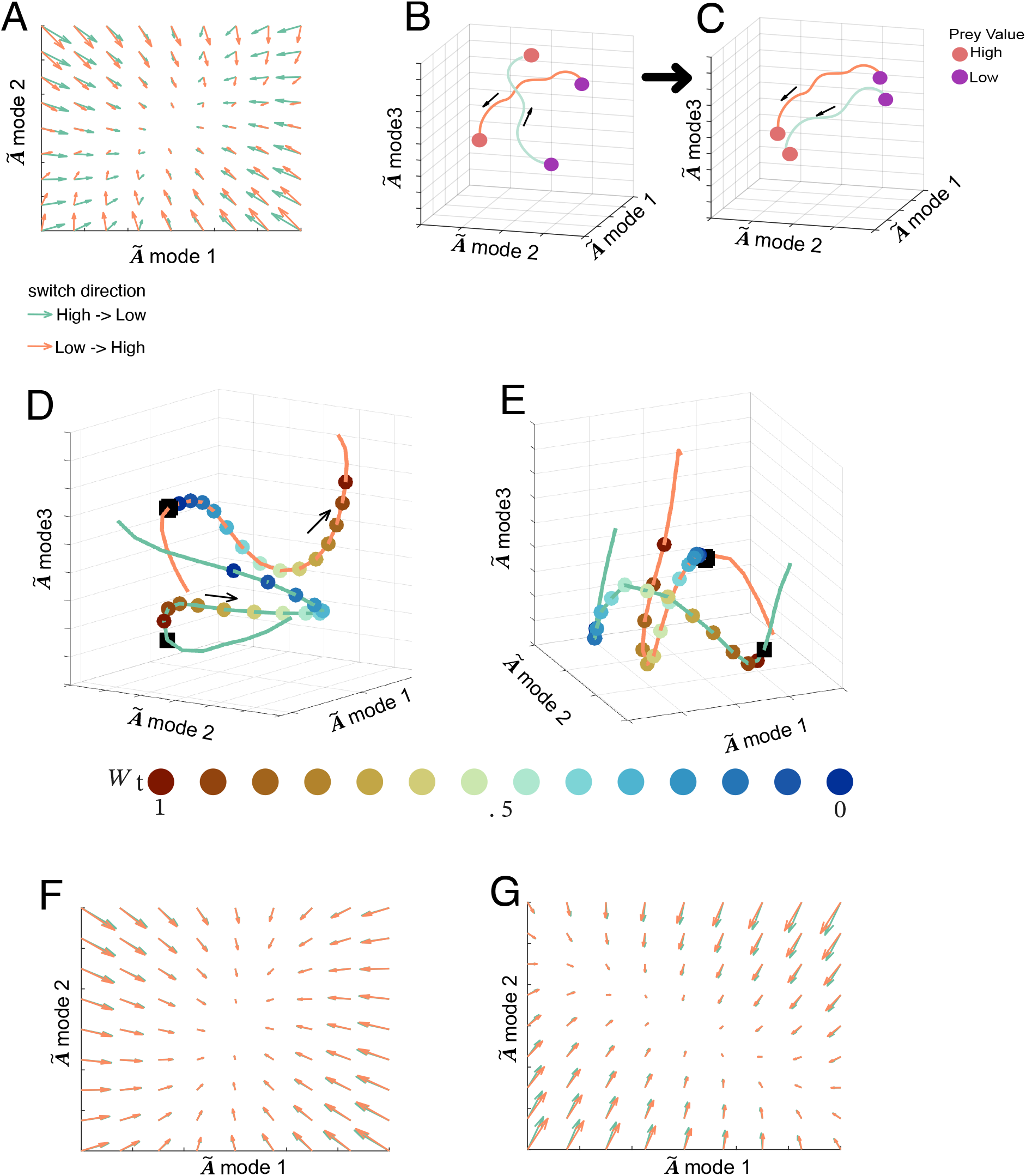
**A**. Arrows show the estimated dynamics vector field, fitted using DMD on the average neural population data (ACC). Vectors represent the derivatives in both directions for each point. Green arrows show high-to-low W_*t*_ switch dynamics. Orange arrows show low-to-high W_*t*_ switch dynamics. The first 2 dimensions of average estimated dynamics are shown (simulated from equation 12, **Methods**). **B** and **C** show a theoretical diagram of the neural state space during switching for a system with separable switch direction conditions (**B)**, and (**C)** that if both conditions share dynamics (ie., topological conjugacy), they will be alignable. **D**. low-dimensional dynamics subspaces defined by DMD on average Neural firing rates in ACC populations. **E**. is the same **D** but for PMd. The colored dots represent different average levels of W_*t*_ during the switch window. **F** (ACC) and **G (**PMd) show vector fields after fitting DMD and applying DSA, showing that the vectors are realigned to match dynamics between switch direction conditions.

Next, we used DSA on all pseudo populations together, forming high-dimensional (d=103 in dACC and d=165 in PMd, both accounting >80% variance) dynamic mode vector fields. Before DSA, comparison of dynamics between switch directions only had a modest correlation distance (dACC: 0.19, PMd: 0.14). After DSA, both correlation distances between dynamics were high (dACC: 0.93, PMd: 0.89), with these alignments occurring with high-fidelity compared to random permutations (z-test on randomized DSA loss permutations: dACC: z = -18.45, *p* < 1x10^-5^; **Figure 4F**; PMd: z = -10.12, *p <* 1x10^-5^, **Figure 4G**). The implication of this dynamic alignment is that neural activity flowing in both switch directions is underpinned by conserved dynamical rules. Namely, it points to a shared set of attractor mechanisms that control the flow of the policy mixing representation W_*t*_. A key advantage of a conserved attractor for W_*t*_ is it can be repurposed in novel environments rather than relearned de novo (Khona and Fiete, 2022). A reused neural W_*t*_ control allows immediate compositionality of low-level controllers, even when environmental contingencies change (e.g., prey value or speed).

## DISCUSSION

Reward-based choice in continuous contexts poses important and parallel low-level control and meta-control problems (Yoo et al., 2021). Solving these hierarchical problems can be quite complex; the solutions require having low-level policies that dictate the particular controls towards specific target rewards, as well as high-level meta-control policies that dictate how low levels should be combined at any moment. Here we find that macaques can solve this problem during an active pursuit task involving multiple prey, and that their behavior is parsimoniously explained as the result of the deployment of a signal that continuously controls a mixture of low-level control policies. The implementation of a continuous meta-control is necessary. Previous work, using approaches like HMMs, has focused on discrete behavioral states that cannot even in principle be combined (for example, forward and backward motion). However, discrete strategies are a special case of degenerate meta-control. In the more general case, goals such as multi-agent pursuit can be composed up multiple strategies. Using our control theory approach, we found a representation of our posited meta-control signal W_*t*_, to be reified in population computations. One major advantage of such a signal is that it can serve as a repurposed dynamics in novel environments (Driscoll et al., 2022; Khona and Fiete, 2022), allowing multiple pursuit control policies for novel objects to be rapidly composed together.

Our approach represents a shift in perspective from traditional approaches to ethogramming, which primarily rely on minimally constrained, unsupervised learning to identify states (Anderson and Perona, 2014; Nair et al., 2023; Pereira et al., 2020; Voloh et al., 2023); therefore, those methods remain agnostic to the agent’s latent cognitive processes. Instead, we found that uncovering the moment-by-moment subject choice requires positing task-driven constraints on how they subjects solve the problem (Kwon et al., 2020). Here, that is their policy mixing (W_*t*_) and controller policies over different prey goals. In addition, the finding that our continuous controller model explained subject control substantially better than a two-state HMM (comparisons shown in **Table S1, Figure S2-S3**) further supports the engagement of continuous compositional control over discrete goal-pursuit. Such a finding indicates that understanding ethological, high-dimensional behavior must go beyond simply categorizing discrete behavioral motifs.

The pursuit of fleeing prey and its inverse, evasion of pursuing predators, is a behavior that is observed across the animal kingdom, from insects to primates (Cooper and Blumstein, 2015). We show that pursuit choices between multiple prey can be well understood from a control-theoretic approach. More generally, that approach opens the door to understanding a wide range of naturalistic behaviors not well modeled by standard approaches (Pearson et al., 2014). These include, for example, cooperative hunting, intraspecific agonistic behaviors such as threatening and fighting, and a variety of mating behaviors. More generally these models can help us understand any situation in which changes of mind can occur on-line, in the midst of continuous action (Yoo et al, 2021). We anticipate that our approach, then, can help neuroscientists begin to understand these types of continuous, embedded, and ultimately, naturalistic decisions.

## METHODS

All animal procedures were approved by the University Committee on Animal Resources at the University of Rochester or the Institutional Animal Care and Use Committee at the University of Minnesota. All experiments were conducted, and designed to comply with the Public Health Service’s Guide for the Care and Use of Animals.

### Subjects

Two (males) rhesus macaques (Macaca mulatta) served as subjects. Previous training history for these subjects included a variant of a Wisconsin card-sorting task (subject H, Sleezer et al., 2016), basic decision-making tasks (subject H, Strait et al., 2014; Fine et al., 2023), and two foraging tasks (both subjects, Blanchard et al., 2014A and B).

### Electrophysiological recording

One subject was implanted with multiple floating microelectrode arrays (FMAs, Microprobes for Life Sciences, Gaithersburg, MD) in the dACC and PMd. Each FMA had 32 electrodes (impedance 0.5 MOhm, 70% Pt, 30% Ir) of various lengths to reach dACC. Neurons from another subject were recorded with laminar V-probe (Plexon, Inc, Dallas, TX) that had 24 contact points with 150 μm inter-contact distance. Continuous, wideband neural signals were amplified, digitized at 40 kHz and stored using the Grapevine Data Acquisition System (Ripple, Inc., Salt Lake City, UT). Spike sorting was done manually offline (Plexon Offline Sorter). Spike sorting was performed blind to any experimental conditions to avoid bias.

### Experimental apparatus

To control their computer avatar, subjects used a joystick that was a modified version of a commercially available joystick with a built-in potentiometer (Logitech Extreme Pro 3D). The control bar was removed and replaced with a control stick (15 cm, plastic dowel) topped with a 2″ plastic sphere, which was custom designed to be ergonomically easy for macaques to manipulate. The joystick position was read out in MATLAB running on the stimulus control computer.

### Task description

At trial start, two or three shapes appeared on a gray computer monitor placed directly in front of the subject. The yellow (subject K) or purple (subjects H) circle (15-pixels in diameter) was an avatar that represented the subject, and began at the center of the screen. Subject position was determined by the joystick and was limited by the screen boundaries. A square shape (30 pixels in length) represented the prey(s). Prey movement was determined by a simple algorithm (see below). Each trial ended with either the successful capture of the prey or after 20 seconds, whichever came first.

Successful capture was defined as any spatial tangency between the avatar circle and the prey square. Capture resulted in an immediate juice reward, whereby the juice amount corresponded to prey color as follows: 0.3 ml for orange; 0.4 ml for blue; 0.5 ml for green; 0.6 ml for violet; and 0.7 ml for cyan.

Prey movement was generated interactively using an A-star algorithm. Specifically, for every frame (16.67 ms), we computed the cost of 15 possible future positions the prey could move to in the next time-step. These 15 positions were equally spaced on the circumference of a circle centered on the current position of the prey, with a radius equal to the maximum distance the prey could travel within one time-step. The cost was based on two factors: the position in the field and the position of the subject’s avatar. The field that the prey moved in had a built-in bias for cost, which made the prey more likely to move toward the center. The cost due to distance from the avatar of the subject was transformed using a sigmoidal function: the cost became zero beyond a certain distance so that the prey did not move, and it became greater as distance from the avatar of the subject decreased. From these 15 positional costs, the position with the lowest cost was selected for the next movement. If the next movement was beyond the screen range, then the position with the next lowest cost was selected until it was within the screen range.

The maximum subject speed was 23 pixels per frame (each frame = 16.67 ms). The maximum and minimum speeds of the prey varied across subjects and were set by the experimenter to obtain a large number of trials. Specifically, speeds were selected so that subjects could capture prey on no more than 85% of trials. To ensure sufficient time of pursuit, the minimum distance between the initial position of each subject avatar and prey was 400 pixels.

### Behavioral analysis

#### Estimating contribution of each target to control and switching

The goals of this study were multi-fold: determining how subjects weigh the contribution of each target to their pursuit choices at a given moment, and assaying how neurons encode this target weighting and change-of-mind. If we recover this target weighting, we can address the core question of how neural populations encode this relative target weighting and choice of switching between targets.

To do so, we developed a method for decomposing the subject’s instantaneous behavioral movement control (i.e., velocity) into how it is driven by two different targets that are being pursued. There are several approaches one could take to solving this problem. This problem requires solving a latent variable problem. The latent variable here corresponds to the relative weight (*W*_*t*_) applied to subject control (*U*_*t*_).

To estimate the latent *W*_*t*_ (we assume or explicitly state time dependency, moving forward), several approaches could be taken. All of them require assuming some form of generative model assumption. Namely, we must postulate *a priori* how *W*_*t*_ are generated according to the subject. Because *W*_*t*_ is continuous in a bounded range [0,1] and in time, we limited our choice of method to those that approximate the dynamics of *W*_*t*_. But, because we wanted to estimate *W*_*t*_ rather than model *W*_*t*_ dynamics – (the latter is achievable with dynamic systems models or function approximators) – we can model the whole time-series of *W*_*t*_ within a single-trial using radial basis functions. In addition, because we model the time dependency of *W*_*t*_, the locality of radial basis functions (RBFs) offers several advantages. RBF approximations to *W*_*t*_ allow optimizing a smooth trajectory, sparse representations, and it can model complex and nonlinear changes in control dynamics (Bishop, 2006). Using RBFs for approximation of complex target weighting dynamics means we can estimate the *W*_*t*_ dynamics without the restrictions of linear dynamics systems, or assuming the types of inputs into a dynamic system. From a model-fitting perspective, the only regularized assumptions we make is smoothness between adjacent RBFs (graph Laplacian prior).

Algorithmically, we solve this problem using maximum-likelihood. The maximization step is performed numerically due to the nonlinear relation between model parameters and the loss function. Optimization was performed using Bayesian adaptive direct search (BADS; Aceribi & Ma, 2017) The collection of model parameters θ optimized per-trial includes RBF weights (*w*_*i*_), a shared width parameter for RBFs, and R for each linear-quadratic regulator (LQR) controller (Q is set to 1) to compute the controller gains (K), and the joystick state dynamics observation noise *ϵ*∼*N*(0, σ). Our aim is to solve this mixture of controllers problem by computing the summed log-likelihood (equation 1) for subject control versus the model-predicted:

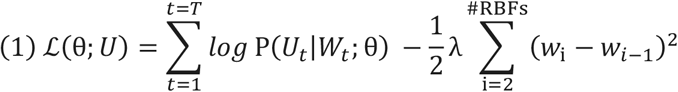

Equation 1 represents the optimization loss function. The first term in Equation 1 is the log-likelihood and the right hand side is a Laplacian smoothing regularization between RBF amplitude weights (*w*_*i*_). The likelihood of the observed cartesian control signal (*U*_*t*_: *observed* joystick velocity in x and y dimensions), given the predicted controls for each target 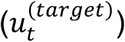 at each time-step is denoted as:

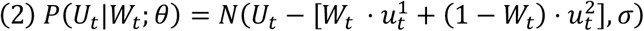

To optimize this model formulation, and at each optimization iteration for a given trial, we compute *W*_*t*_. The function approximation is computed using the optimized weights for each RBF (equation 3), which is subsequently range transformed through a sigmoid function [0,1]:

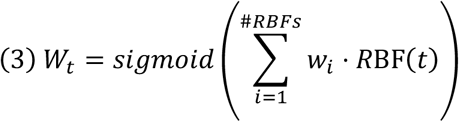

After computing the weighting trajectory, we compute the error between each controller and subject’s cartesian joystick position. Then we obtain the optimal gains (via the discrete algebraic Ricatti equation, dlqr.m in Matlab) from each infinite-horizon LQR, corresponding control signals for each target (equation 4), and weight them accordingly (equation 5) to create a predicted subject’s control signal:

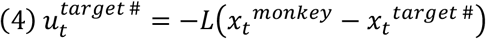

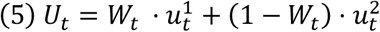

#### Converting model fits over N-bases and lambda into predictions

After fitting a series of weights for RBFs to each individual trial over a series N-bases (11, 21, 26, 33) and Laplacian regularization weights (1x10^-4^, 1x10^-3^, 1x10^-2^), we average the predicted *W*_*t*_ trajectory from each fit rather than select a best. This was done because we wanted to obtain the most likely trajectory over the space of possible fits. Model averaging reduces overall uncertainty in *W*_*t*_ and helps mitigate issues with over-fitting (Murphy, 2022). To average, we use an approximation to Bayesian model averaging that employs the Bayesian information criterion (BIC) to compute weights for each model parameterization fit to each trial; because the BIC is a Laplace approximation to the model evidence, this approach approximates Bayesian model averaging when a uniform prior is used (Hoeting et al., 1999). BIC model-averaging weights were created (equation 6) to derive a pseudo posterior probability for each fit per trial. These weights were used for each trajectory.

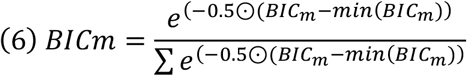

#### Finding behavioral switch points

To separate instantaneous moments into pursuit versus change-of-mind points between targets, we first searched each trial to find points where *W*_*t*_ =0.5. These *W*_*t*_ points denote crossings between targets. We excluded any putative switch points that occurred within 150 ms of the trial start time, under the assumption that these likely represent a correction to an initial choice rather than true pursuit switching or change-of-mind. Using the remaining potential switch points, we created a window of -300 to +300 ms around these points. The switch trajectories were then categorized as falling into one of three categories. The main category (1) was those where a full switch was made between targets, the other two were either (2) partial switches or (3) were unclassifiable. To achieve this classification, we used a PCA across all switch trajectories from within a session. The 1^st^ principal component (PC) always captured the sigmoidal shaped full switch trajectory of interest. We computed the cosine similarity of this 1^st^ PC score and all switch trajectories and retained those with a similarity of > 0.9. We ran a PCA again on this subset, with the 1^st^ PC again producing the canonical sigmoidal trajectory. In this case, we retained those with cosine similarity of >0.97. This threshold proved to be conservative enough to remove all partial switches, based on visual inspection. In total, an average of 35% of *W*_*t*_ crossing-points were labeled as legitimate switches and forwarded for further analysis. Switch onset and offset timing were determined using a linear fitting change-point algorithm that looked for changes in mean and slope of the *W*_*t*_ trajectory. To find the switch start, we searched backward in time from the crossing of *W*_*t*_ = 0.5. Switch ends were found by searching forward in time.

#### Switch statistics

The factors underlying the propensity to make switches were determined using a Lasso approach to a logistic (binomial) regression. We asked if switch probability was sensitive to the time elapsed in trial, the Δreward of prey, and several prey distance variables. Specifically, we included the mean distance to the prey being pursued and the mean distance of the prey switched to, where the mean is calculated in the 100 ms leading up to a switch. We also included a factor to determine the mean rate of change of distance (distance closure) between both of these preys, too. The distance closure captures whether either prey getting closer or farther away (on average) altered switch probability. To derive an interpretable centering for each predictor in this model, we extracted trials in which there were no switches. Put differently, we wanted to ask whether the time-elapsed in a trial where there was a switch was significantly then the median time-elapsed in trials without a switch. For example, for trials with switches, we centered the time-elapsed in a trial on the median trial length of trials with no switch. Similarly for other variables such as means and change of distances, we centered switch trial variables at the median of the control trials. The control trial value, per trial, computed at the mid-point at each control trial. The median of these was used for centering, making our predictors interpretable with respect to the being greater or less than these controls. The only variable not centered was Δreward.

All of these predictors and their (up to 2 way) interactions created 22 predictors to be modeled with. We used a Lasso (L1 regularized) model with 5 K-fold cross-validation to determine parameter values. Because we wished to model both monkeys together to determine globally relevant parameters, and each monkey differed in total trial numbers, we used a stratified bootstrapping approach. We performed 50 bootstraps, with the model being fit each time, using 1000 stratified, randomly selected trials per monkey for a total of 2000 trials. We determined parameter significance using 99% confidence intervals over the resultant parameters, determining those non-overlapping with zero.

### Neural analysis

#### Single neuron encoding of task variables

Neuron encoding of task variables across all experimental time was estimated using a linear regression on single-neuron firing rates (gaussian filter smoothed at ∼32 ms). The z-scored firing rate vector (***y***) was created by concatenating all trials starting from the estimated reaction time of each trial to the end of target pursuit. We aimed to determine whether single neurons encode the latent target choice weight (*W*_*t*_) either independently or mixed linearly and/or nonlinearly with subject cursor speed and the difference in capture values between the two prey. The regression model for each unit was as follows:

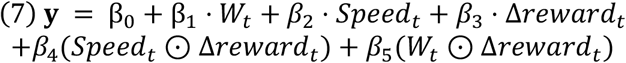

As we were assessing selectivity, and correlations between task variables (e.g., speed and *W*_*t*_) were low (max = 0.2), we fit each model using maximum-likelihood without regularization. Statistical significance was determined by permuting the individual task variables into random time-bins first, creating their interactions, and then fitting the permuted data model. Effects and interactions were considered significant at *p* < 0.05. We controlled the chance level proportion of neurons with significant variables by using a false-discovery rate of 1x10^-5^ across neurons.

#### Demixed PCA and population decoding

To determine how neuronal populations represent a change of mind and the pursuit weight *W*_*t*_, we used demixed principal components analysis (Kobak et al., 2016). This method allows us to determine whether the population has linearly decodable representations of the target pursuit weight. In addition, we can use the same model to learn neural population modes that track change-of-mind over time, independent of the switch direction (from low to high or high to low value target); these are the condition independent modes.

To decompose the data, we first sorted all switch points, during a trial, for a given subject, and within a session, into those that switched from a low to high or high to low value target. Our aim is to find a linearly decodable representation of *W*_*t*_. Under the assumption that *W*_*t*_differs in a mirrored fashion (high to low versus low to high *W*_*t*_), across time, for these switch directions, we can take the decoding subspace as the representation subspace for *W*_*t*_. For example, this means that a representation for *W*_*t*_can be found by finding the decoding hyperplane. To do this, we first formed a firing rate matrix tensor that included soft-range normalized (constant of 5) and grand-mean centered firing rates. The tensor was shaped as N (neurons) x switch direction (2 levels) x time-points (30, [-15,15] around switch point) x trials. All the time-points were centered at the switch point (*W*_*t*_= 0.5), and the firing rates for each trial were time-warped to the median change-of-mind onset and offset points; warping was done using a cubic spline interpolation scheme. For analyses, these firing rate tensors or pseudo-populations were created separately for both dACC (170 neurons from subject H and 25 from subject K) and PMd (330 neurons from subject H).

Using the firing rate tensors, we then used dPCA as a decoder following several steps. First, for each of 1000 repetitions, we randomly sampled (stratified monte-carlo) the pseudo-populations and held-out 5% of trials (balanced between switch directions). A training and test firing rate tensor was created, with each tensor averaged over trials. A dPCA model was then trained using a fixed regularization of 1x10^-5^. The test-set was projected along the top two decoder dimensions for the switch direction. We took the decoded switch direction to be the minimum distance of the projected test data to the projected training data. This classification was computed for each of the 30 time-points in the switch window. Second, we established significance thresholds by repeating the above procedure, but randomly swapping trial labels to create a null decoding accuracy distribution. We created the null distribution by repeating the 1000 iterations 50 times. For each of the 50 sets, we computed the null decoding accuracy. We determined that decoding was statistically significant when the decoding accuracy was greater than the null for 100% of the 50 sets.

#### Dynamics comparison overview

One of our core hypotheses is that a change of mind reflects a certain traversal of *W*_*t*_ by neural activity. The core implication is that this activity evolves with certain dynamics along a topology in a low-dimensional neural state space. One hypothesis this predicts is that traversal of *W*_*t*_ from either a low to high or high to low switch should use similar dynamics. The implication is that the proper way to compare switching conditions for a conserved representation of *W*_*t*_ is in their dynamics, not their static geometry.

We explain the problem differently, here. An issue with comparing two putative dynamics systems to determine alignment is that state space geometry is not equivalent to topological (dynamics) conjugacy. Namely, two distinct systems can have identical (or distinct) geometries yet have different (or similar) dynamics driving them; for example, assuming linear systems, they could have very different eigenvalues that determine the dynamics, despite both being stable systems. A simple example in two-choice decision making is a 2D system driven to either choice by a line attractor, saddle point, or bistable discrete attractors will exhibit the same geometry on average (Ostrow et al., 2023; Galgali et al., 2023). This means that neuronal state spaces dynamic portraits cannot be compared directly using geometric metrics (e.g., canonical correlation, distance measures, etc.), but require comparison in terms of dynamics. Next, we give an overview of the method we use to test this, and then outline the steps for computing them.

### Two-step approach for conjugacy: Dynamic mode decomposition and dynamic similarity

To compare two systems dynamics, we adopted the two-step approach of Ostrow et al. (2023). We first apply a version of dynamic mode decomposition (DMD) with unsupervised control to estimate neural population dynamics during a pursuit switch (Fieseler et al., 2020). DMD is a data-driven approach to dynamical systems estimation for high-dimensional systems (explained below). In this second step, we ask whether the dynamics’ systems operators dictating the flow in neural state-spaces can be aligned. The core idea is that if two linearly approximated dynamical systems exhibit conserved dynamics across conditions, but potentially subject to linear distortions that distinguish their dynamics, they will have matching eigenspectra and be alignable, i.e., topological conjugacy (Budišić et al., 2012).

#### Step 1. Compute dynamics from DMD

Here, we provide a brief overview on DMD as relevant to determining dynamic similarity; thorough overviews are provided in several other places (Kutz et al., 2016). At its core, DMD is a data-driven approach that attempts to linear an infinite-dimensional linear system operator of a finite nonlinear dynamic system. Learning a linear system operator means that linear systems analysis is applicable and allows easy comparison of systems. Because we model as a linear system, the dynamics found by DMD are either growing/decaying exponentials or oscillatory (complex eigenvalues). Using DMD for a linear system that describes evolution of neural activity, it takes the form of a linear, discrete dynamic system:

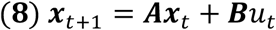

As applied to our neural data, in equation 8, ***x***_*t*_ is the firing rate vector of neurons at time *t*. ***A*** is the linear transition operator with dimension of # neurons x # neurons, ***B*** is the control (*u*) scaling for each unit. Because we wanted to account for potential input differences, but neither ***B*** nor *u* are known a priori, we used a recent DMD method that learns a latent *u* and ***B*** before estimating the dynamic modes (Fieseler et al., 2020). This approach aims to estimate sparse latent neural control (*u*_*t*_) signals that resolve event-changes, akin to a switching-linear dynamical system. Next, an approach known as exact DMD is applied to estimate ***A* (**Kutz et al., 2016). Because DMD is a dimensionality reduction approach, too, exact DMD allows us work directly in the lower-dimensional space spanned by the number of dynamic modes. Specifically, we directly estimate a lower-dimensional transition operator with rank equal to the number of dynamic modes (***r***),***Ã***_**r x r**_.***Ã*** has dynamics described by eigenvalues shared with the full matrix ***A***, but ***Ã*** operates on the low-dimensional dynamics rather than the neuron space of ***A***. We want ***Ã*** because we will compute it for each switch direction conditions, and align them with a dynamic similarity metric. Exact DMD computes ***Ã*** in several steps, first by splitting data ***x*** into ***X*** = (***X***: *t* ∈ *T* − 1) and ***X***’ = (***X***’: *t* + 1 ∈ *T*), with *T* equal to the time-series length. ***Ã*** is now computed as follows, where any item with *∼* is truncated to ***r*** modes:

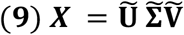

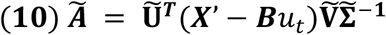

Equation 9 is the singular value decomposition of ***X***, with ***r*** truncated left-(and right) singular-vectors and singular-values. In eq.10, we see that we can directly project ***A*** into the lower dimensional space of ***Ã*** that defines the dynamics along the retained modes. Therefore, ***Ã*** defines the estimated systems dynamics equivalent to fitting the latent state transition operator in linear dynamic systems. For predicting the fitted dynamics, we can equally project an input matrix ***B*** into 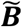 (equation 11) for simulating the learnt dynamics (equation 12, Figure 4D-E; full derivation is found in Proctor et al., 2016).

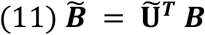

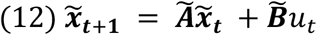

#### Dmd parameterization and estimation for average and all trials

We now describe the specific parameterization used for the neural switching window data. The main choice for DMD analysis is the rank **r** (number of modes). We followed a standard approach by using the first *n* modes from the SVD of ***X*** that covered 80% variance (estimated from 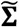). When estimating this for the average neural population trajectory, for both switch conditions, we found 10 DMD modes for both ACC and PMd; this DMD shows they were indeed low-dimensional and were used for plotting average dynamics state spaces in Figure 4 of the main text.

For full similarity analysis (DSA) to compare the dynamics ***Ã*** between neural switch direction conditions, we did a DMD on a pseudopopulations of trials (similar to Ostrow et al., 2023) for each ***Ã***. This is important for two reasons. First, because the average dynamics can look falsely equivalent without accounting for variability (Galgali et al., 2023). Second, we can estimate a singular ***Ã*** that accounts for expected dynamics across pseudopopulations; this approach is similar to concatenation of conditions in PCA for neural state space analysis.

To make the pseudopopulations, we first estimated the maximum number of trials, per switch condition, per neuron. Taking 15% of this number (∼5 trials), we then made subsampled (no replacement) pseudopopulation trials of firing rates. The sampled trials per neuron were averaged within each pseudo-, and all neurons firing rates were stacked into the matrix ***X***. This process was repeated 100x for each switch direction. For ACC, we estimated 103 modes, and PMd, we estimated 165 modes. Note, the number of modes is substantially higher here than for the average system because we are accounting for variability in dynamics across pseudo trials. Finally, the two separate matrices ***X*** for the high-to-low *W*_*t*_ switch and low-to-high *W*_*t*_ switch are both submitted to the DMD pipeline to retrieve ***Ã*** for each.

#### Step 2. Dynamic similarity analysis

To compute the similarity between dynamics of both switch conditions (***Ã***), we use the proper derived metric from Ostrow et al. (2023). We will denote the high-to-low ***Ã*** as ***Ã***_***H***−>***L***_ and the low-to-high condition ***Ã*** as ***Ã***_***L***−*>****H***_. In short, when comparing the similarity in dynamics, we want to compare their vector fields defined by ***Ã***. As noted by Ostrow et al. (2023), when attempting to compare the alignable similarity ***Ã***_***H***−>***L***_ to ***Ã***_***L***−>***H***_ with a geometric (e.g., procrustes rotation with orthogonal matrix ***C***), the matrix ***C*** will only rotate the flow-field vectors. The impact of only rotating vectors is that it distorts dynamics structure in the vector field. Instead, Ostrow et al. (2023) propose a proper metric they call DSA that preserves dynamics during alignment, by changing vector positions and their rotation, using an orthogonal matrix ***C***:

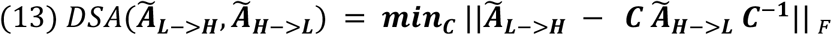

We solve this problem (equation 13) following Ostrow et al. (2023). DSA was estimated by first computing the ***Ã*** for both switch directions from the full data (subsampled). The orthogonal matrix ***C*** was solved for using gradient-descent with Adam (learning rate = 0.01) in PyTorch. Optimization was run for 1500 iterations, but tended to converge in ∼500 iterations.

We also created a permutation scheme to determine chance (noise) level alignment. An orthogonal random permutation of an ***Ã*** destroys the dynamics by changing the eigenvalues. For each DSA permutation (100 total), we initialize a random-orthogonal matrix ***C***. This random ***C*** was then used as a transform against one of the ***Ã*** (equation 13). Then we ran DSA and computed the final loss. Alignment significance was computed using a z-test with the main DSA and the permutations. The theoretical basis for this permutation: if two systems are alignable by shared linear approximated dynamics, then they have common eigenvalues. This is because linear systems’ dynamics are determined by their eigenvalues.

## Supplementary Material

### S1. Mixture of controllers model fitting accuracy on ground-truth recovery

#### S1.1: Recovery of LQR parameters on single prey trials: Q/R ratio is identifiable

In building the optimization procedure for a mixture of controllers, we first asked whether when simulating a single prey trial with an LQR feedback controller, if we could recover the Q/R parameter ratio. To test this, we took prey motion from 10 random trials of subject H data randomly across the 5 sessions. For each trial, we simulated the subject position and control velocity using an LQR controller design with 1 of 5 Q/R ratios; we fixed Q =1. The five Q/R ratios [0.1, 0.5,1.25,1.5, 2.5] arbitrarily place less cost on control as the value goes up; we could have obtained the same controller (and Q/R ratios) by fixing R and changing Q. In the first set of simulations, we include no noise in the simulated controller. In another set we included normal unit process noise. As shown below in Figure S1 (shows mean predicted and estimated), we had near perfect Q/R recovery without noise (98% R^2^) and similar performance with noise (95% R^2^).

**Figure S1.**
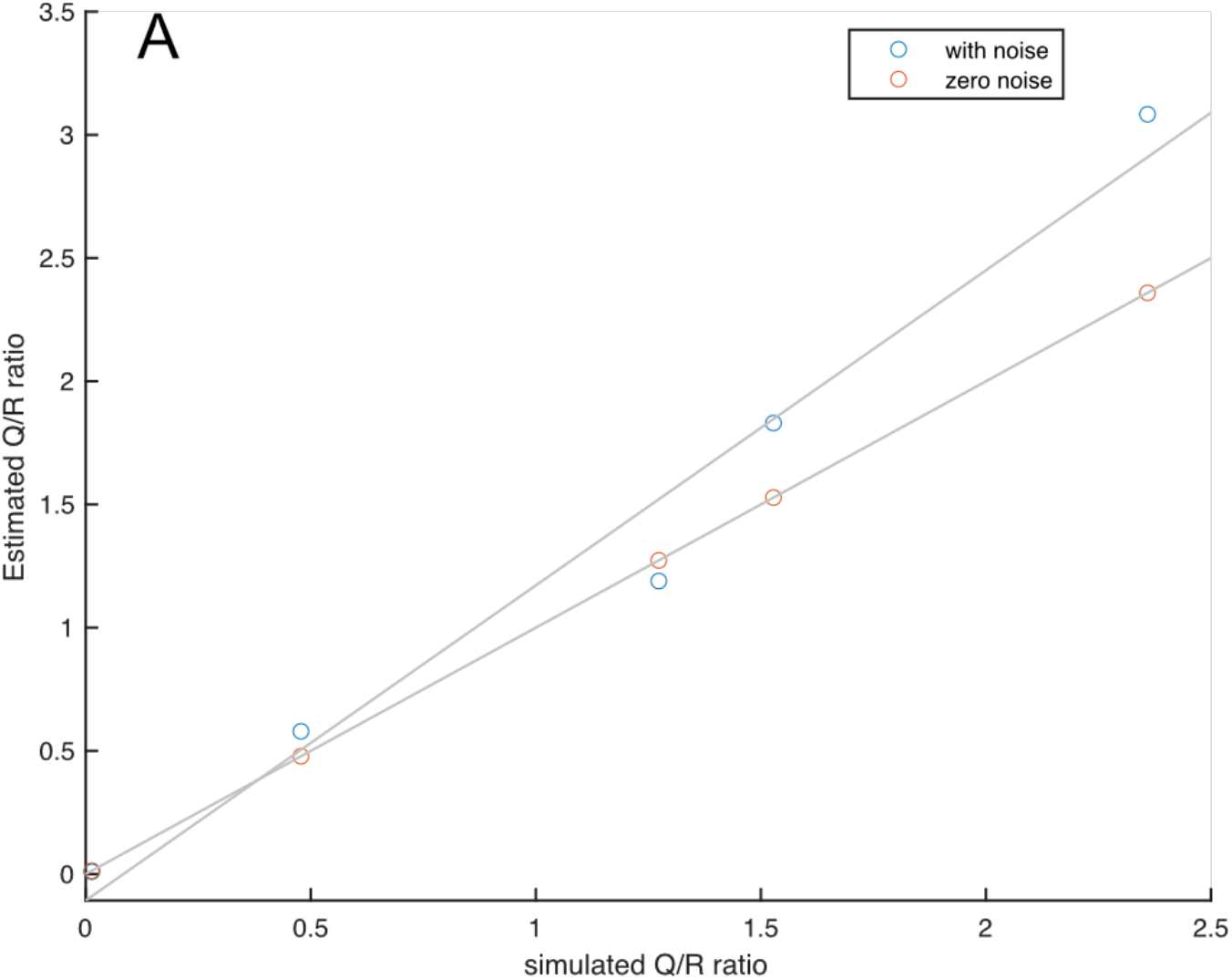
Each point represents the average Estimated Q/R controller ratio from optimizing on simulated single-prey trials. The x-axis indicates the simulated Q/R ratio and the y-axis the estimated. Regression lines of best fit are shown for each. Orange circles indicate simulated optimizations without process noise, and blue circles represent simulations with unit normal process noise.

#### S1.2 Simulated accuracy of model-based control decomposition to W_t_

To test the validity of our modeling approach using basis functions and LQR controllers, we simulated ground-truth data with a known W_*t*_ that we designed. To cover the space from simple to complex W_*t*_ profiles, we considered five different types of W_*t*_. We show examples of these five types in Table S1. To simulate data, we sampled 20 random trials across both subjects’ sessions, with 5 of each having an approximate length of 0.91 sec, 1.45 sec, 2.83 sec, 4.1 sec long. In the main text, we fit across a range of number of bases and model-average. We chose a fixed number of bases in simulation to fix the complexity of the model for comparison to other approaches (e.g., HMM), as increasing basis numbers will increase fit quality. In addition, we also tested model recovery for distortions of the LQR control cost for one of the prey LQRs. Specifically, we considered a simulation when it was given the correct *R* (control cost) (Figure S2, column 1), when the initial *R* was incorrect for the high-value target (Figure S2, column 2), or when the initial *R* was incorrect for the low-value target (Figure S2, column 3). We assessed model fit quality using 3 ways, with our main focus being (1) correlation between true Wt and model-recovered, (2) MSE, and (3) AIC to account for complexity comparison between basis model and HMM (see below). We focus on the correlation here, because the main goal was accurate recovery of Wt. As we show below, even with incorrect specification, our model recovers the true Wt with high-fidelity (Table S1). Additionally, an example recovery applied to a single trial of real subject H data is shown in Figure S1 P-R.

#### S1.3 *Comparative methods for finding Wt:* (1) algebraic and (2) HMM

As a baseline comparison to our model-based approach, we also tested for Wt recovery using either a linear algebraic approach or an HMM. We used the same simulated position, velocity and Wt time-series as used for testing our above approach. For the algebraic approach, we decomposed the simulated instantaneous velocity into two orthogonal components. Specifically, we projected the subject velocity onto each prey and took their orthogonal complement of those vectors. These orthogonal vectors were then projected onto the opposite prey to obtain a measure of independent subject control towards either prey (**V**). These vectors represent the scalar projections of the orthogonal complement vectors (**V’**). The **V’** for each target created a Wt variant as the difference between V for each prey, divided by their sum of both |V|, and normalized it from 0-1. Fits for this approach are in Table S1 (3rd column), and example recovered Wts are shown in Figure S3 first column.

**Figure S2.**
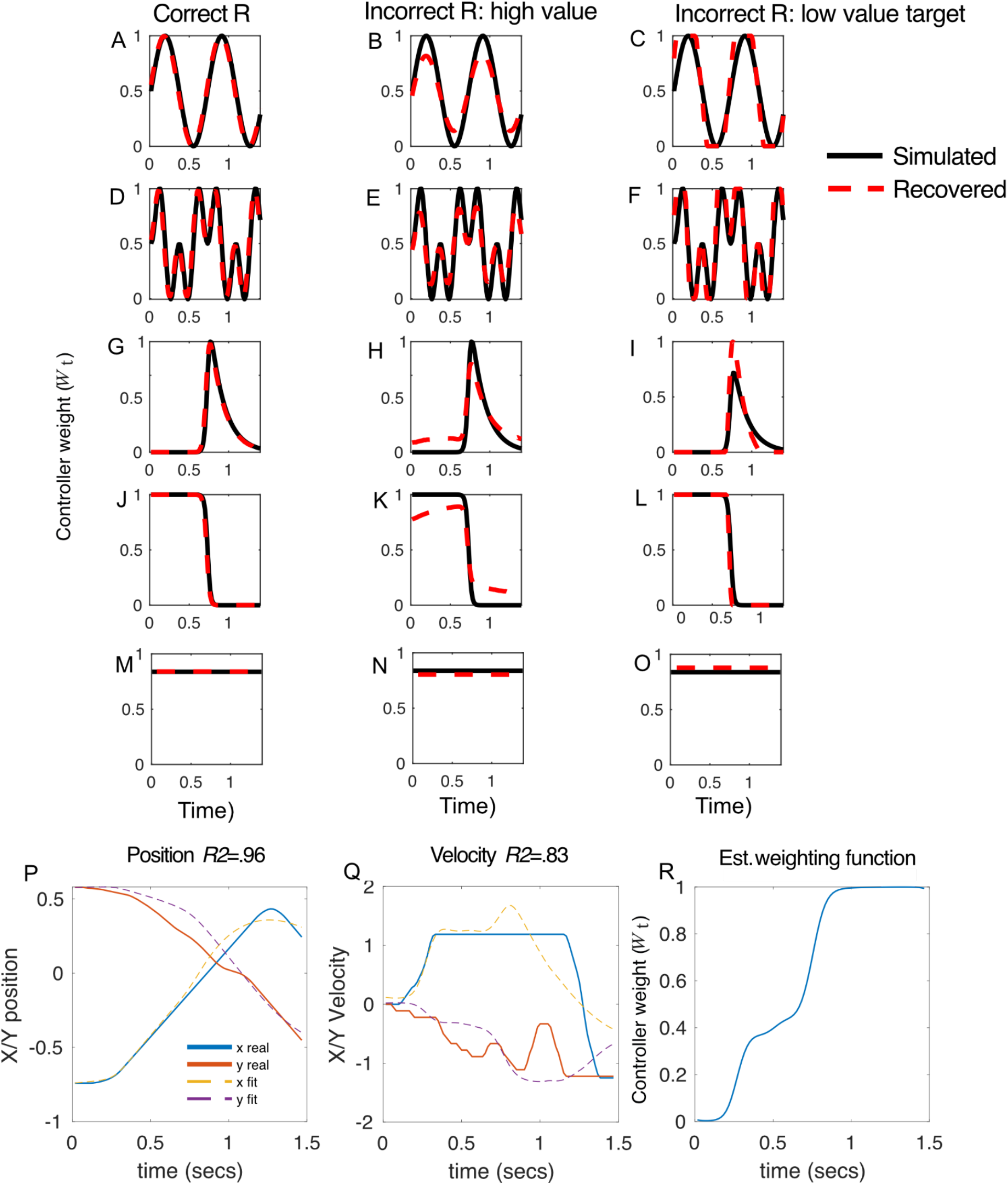
Example outputs from the new optimal -control basis model applied to two-prey trials. Plots A-O show example simulated *W*_*t*_ functions (black line) superimposed with the model-recovered fit (red dashed line). Each row is a different switching *W*_*t*_ function. The first column shows simulated recoveries when the optimization was given the correct R parameters for both LQR controllers. The second and third column show recoveries for when the initial R was mis-specified (but learnt) for either the higher value or lower value prey. In all scenarios, the model performs very well at recovering the ground-truth function (see Table S1). Plots P-R show the model’s fit on a single-trial of real data from subject H. P shows the recovered position using the fit LQRs, Q the velocity traces (control signal), and R the estimated weighting function.

**Figure S3.**
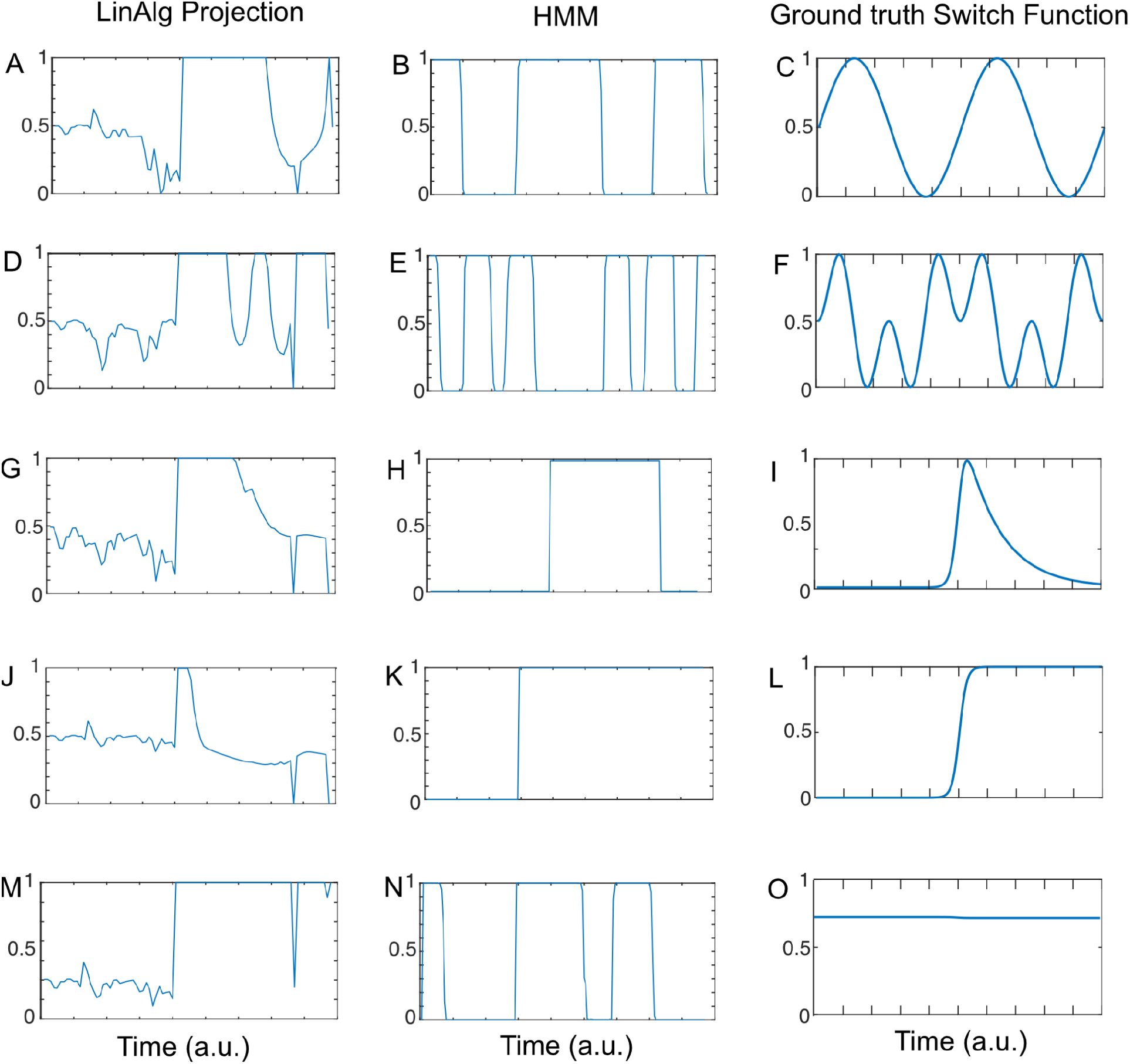
Example predicted *W*_*t*_ for each of the two baseline methods: algebraic projection (column 1: **A, D**,**G, J, M**), and the HMM (column 2: **B, E**,**H, K, N**), for each of the five ground truth switching (column 3: **C, F, I, L, O**) functions used in the simulation study.

The model-based baseline we used was an HMM type called a switching linear regression (Dynamax toolbox). This approach is much closer to modeling the putative impact of inputs to on-going subject control and state transitions (target pursuit). As in our control model approach, we can use the instantaneous subject and prey position discrepancy (**x**\*) as an input signal (i.e., positional error). Then, we can use a two-state HMM (indexed by *k*, one for each prey), each as a separate linear regression that dictates the impact of (X,Y dimension) positional error on subject control. Here, we show this works by rewriting a standard discrete state space model with input:

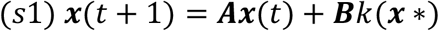

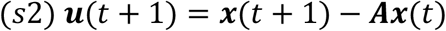

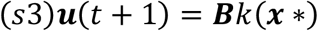

In these equations (*s*1 − *s*3), we show that we can treat transition operator **A** as implicit when we directly model the control **u**, and we only need to learn the linear regression or input parameters (**B)** target error (**x\***) parameters for each HMM state (*k*). **B** is of size 4 (x,y for each of two targets) x 2 (possible HMM states). However, this approach is limited to returning a discretized signal of following a specific target; it is either *k =* 1 or *k* = 2. And more problematically, it can give a good fit to the behavioral control signals while drastically misrepresenting the latent *W*_*t*_ signal; this can occur even when that *W*_*t*_ signal is discretely switching between two states. For a two-state HMM of this type, there are several parameters, here listed with their sizes in subscripts that sum to the total number of parameters (used to compute AIC): (1) *T*_**2 x 2**_: state transition matrix, (2) **π**_**1**_: initial state probability, (3) **B**_**4 x 2 x 2**_: regression weights, (4) **b**_**2 x 2**_: regression intercepts, (5) ***Σ***_**2 x 2**_: (co-)variance matrices. We used the HMM posterior state probability as an estimate of *W*_*t*_ to put the HMM on closer comparison to the basis model output. Fits for this approach are in Table S1 (2nd column), and examples recovered *W*_*t*_ are shown in Figure S3 first column.

## Acknowledgements

We thank Paul Schrater, Xaq Pitkow, Jon Pearson, and Scott Linderman for invaluable discussions and insights on solving the controller problems. We also thank Jeff Johnston for discussion on neural analysis. This project was supported by NIH grants to BYH (DA038615 and MH125377) and by the McNair foundation.

---

**Table S1.**
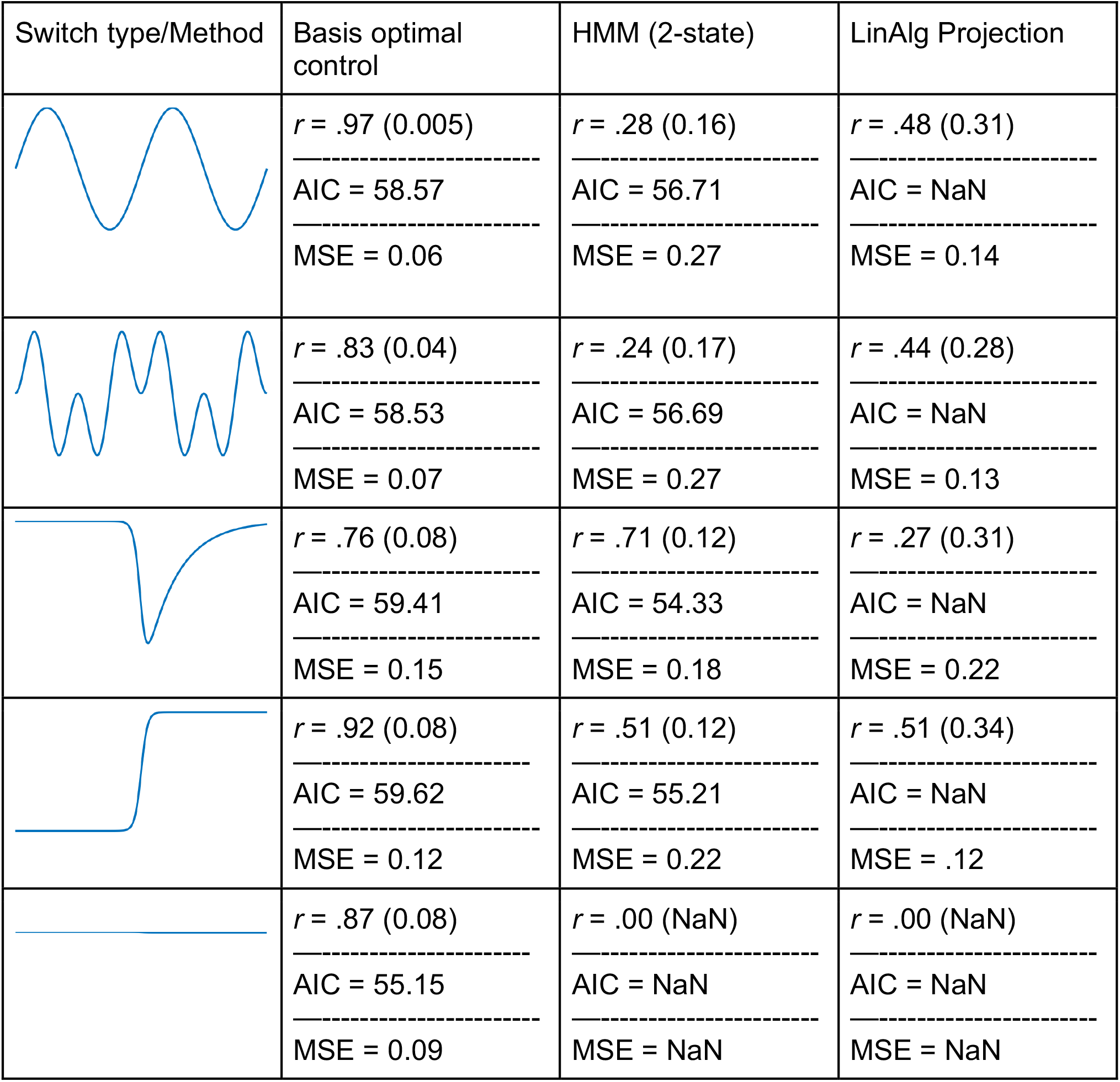
Each method’s correlation (*r*) with ground-truth *W*_*t*_, MSE, and AIC. The first column shows the ground-truth switch function used for simulation and fitting. The first results columns show the mean results for our Basis optimal control model. The HMM and Scalar Linear projection methods are shown in the other columns. Examining both *r* and MSE, we see that our model performs substantially better than all the others, except for 1 instance. For the 3rd switch type (third row), we see the optimal control and HMM perform nearly identical. Correlation *r* standard deviations are shown in parentheses.

| Switch type/Method | Basis optimal control | HMM (2-state) | LinAlg Projection |
| --- | --- | --- | --- |
| | $r = .97$ (0.005)<br>-----<br>AIC = 58.57<br>-----<br>MSE = 0.06 | $r = .28$ (0.16)<br>-----<br>AIC = 56.71<br>-----<br>MSE = 0.27 | $r = .48$ (0.31)<br>-----<br>AIC = NaN<br>-----<br>MSE = 0.14 |
| | $r = .83$ (0.04)<br>-----<br>AIC = 58.53<br>-----<br>MSE = 0.07 | $r = .24$ (0.17)<br>-----<br>AIC = 56.69<br>-----<br>MSE = 0.27 | $r = .44$ (0.28)<br>-----<br>AIC = NaN<br>-----<br>MSE = 0.13 |
| | $r = .76$ (0.08)<br>-----<br>AIC = 59.41<br>-----<br>MSE = 0.15 | $r = .71$ (0.12)<br>-----<br>AIC = 54.33<br>-----<br>MSE = 0.18 | $r = .27$ (0.31)<br>-----<br>AIC = NaN<br>-----<br>MSE = 0.22 |
| | $r = .92$ (0.08)<br>-----<br>AIC = 59.62<br>-----<br>MSE = 0.12 | $r = .51$ (0.12)<br>-----<br>AIC = 55.21<br>-----<br>MSE = 0.22 | $r = .51$ (0.34)<br>-----<br>AIC = NaN<br>-----<br>MSE = .12 |
| | $r = .87$ (0.08)<br>-----<br>AIC = 55.15<br>-----<br>MSE = 0.09 | $r = .00$ (NaN)<br>-----<br>AIC = NaN<br>-----<br>MSE = NaN | $r = .00$ (NaN)<br>-----<br>AIC = NaN<br>-----<br>MSE = NaN |

**Figure S4.**
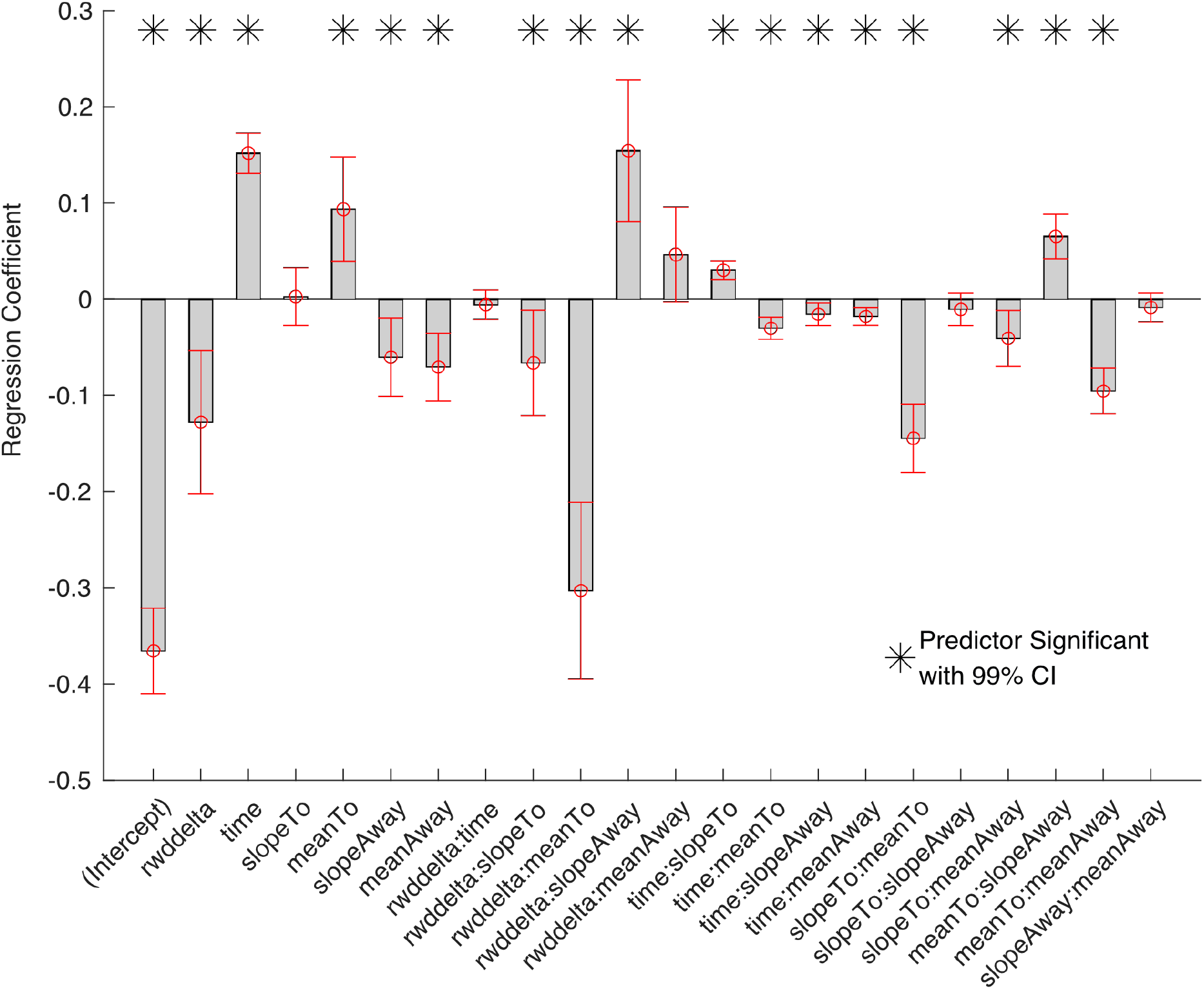
Full results of the Lasso logistic model used to predict the probability of switching P(switch). The coefficients represent the regression parameter for predicting the log-odds. Variables with ‘:’ indicate two-way interaction terms. Below, we list a table of what each of these parameters refers to (Table S2).

**Table S2.**
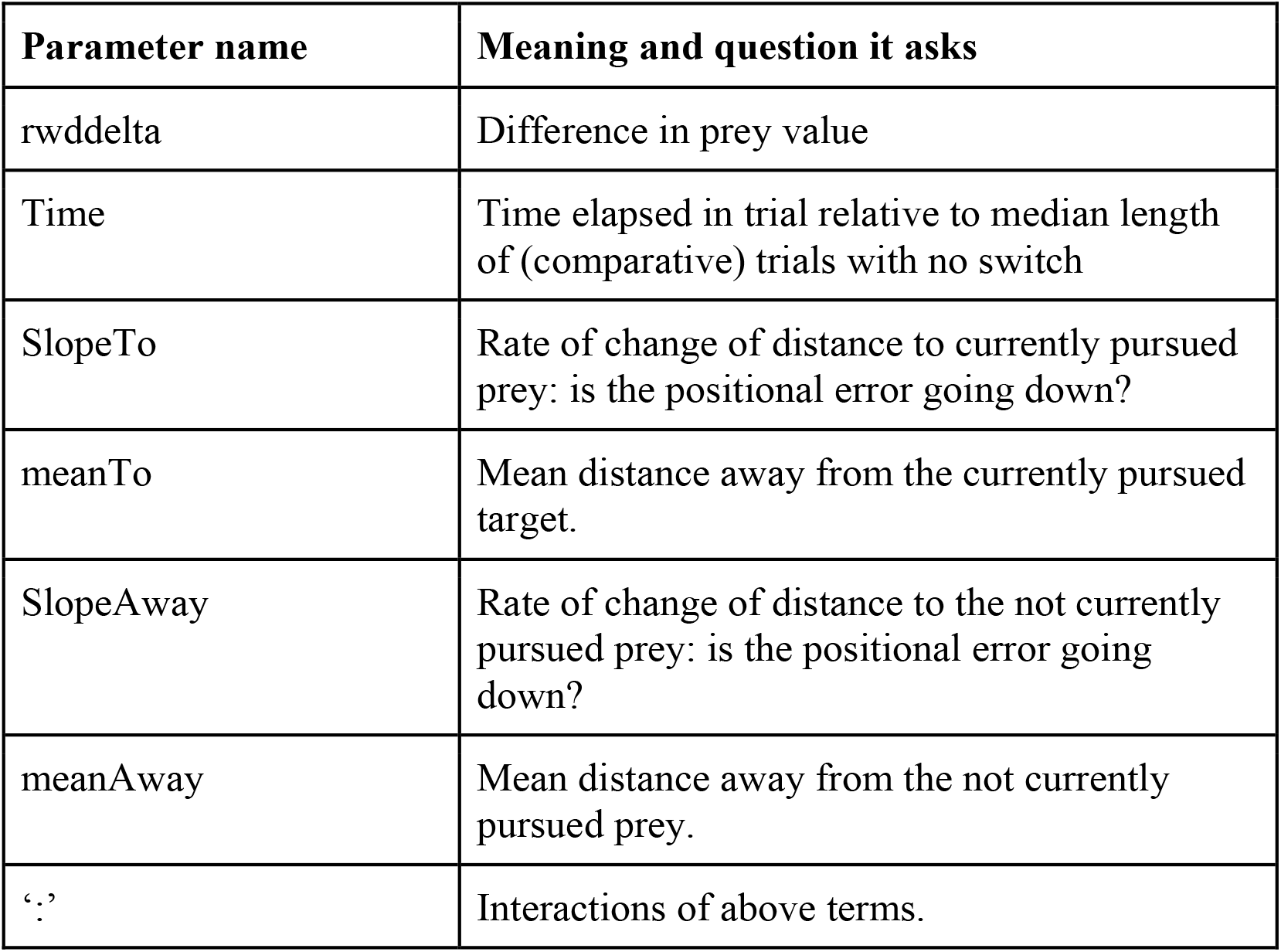
List of all parameters computed as possible predictors for the Lasso logistic regression to analyze probability of switching (P(switch)).

| Parameter name | Meaning and question it asks |
| --- | --- |
| rwddelta | Difference in prey value |
| Time | Time elapsed in trial relative to median length of (comparative) trials with no switch |
| SlopeTo | Rate of change of distance to currently pursued prey: is the positional error going down? |
| meanTo | Mean distance away from the currently pursued target. |
| SlopeAway | Rate of change of distance to the not currently pursued prey: is the positional error going down? |
| meanAway | Mean distance away from the not currently pursued prey. |
| ‘:’ | Interactions of above terms. |

**Figure S5.**
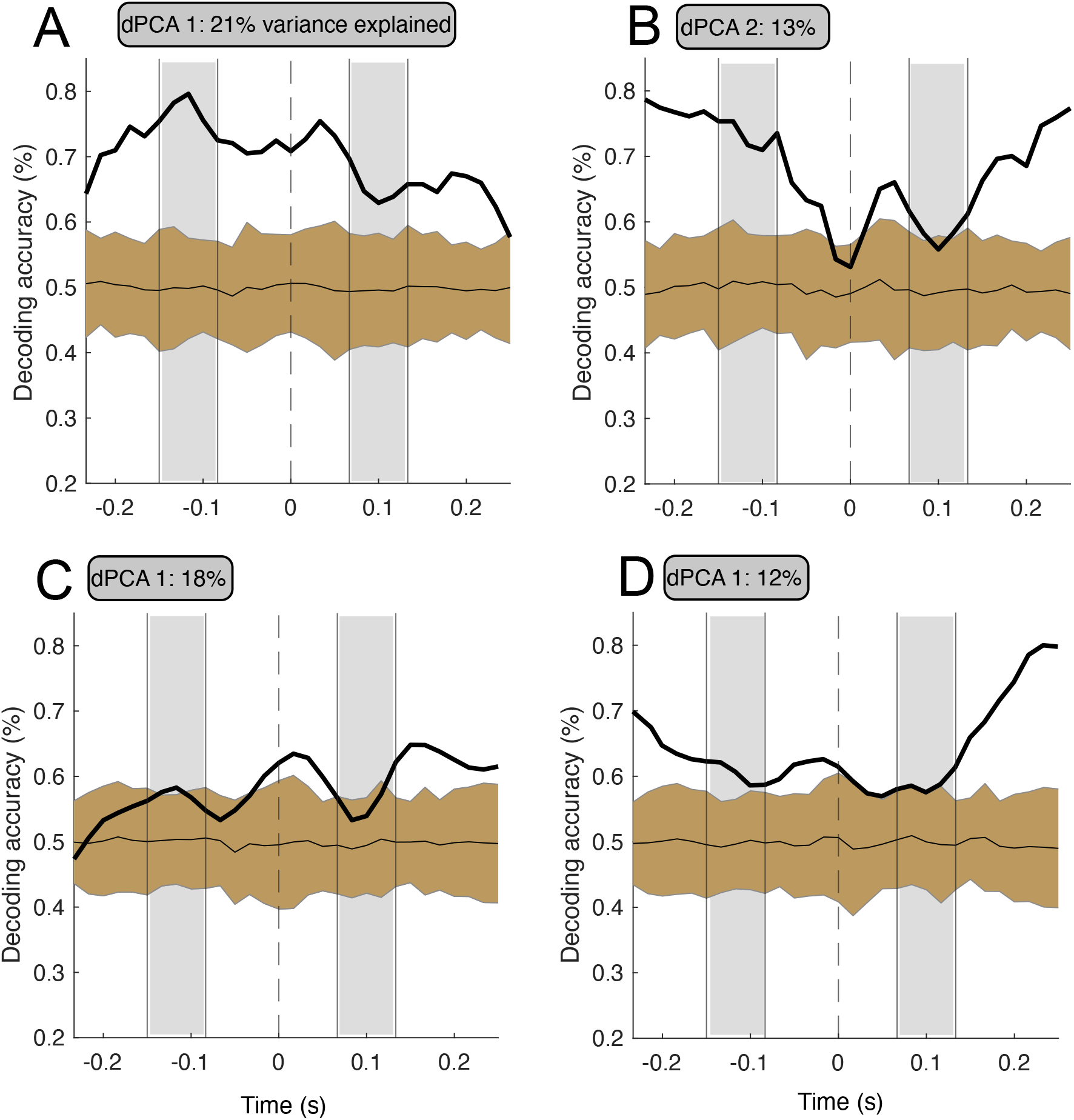
Decoding accuracy for the 1st and 2nd demixed principal components (dPCA) for the switch direction factor in the main text (brown shaded region represents 99% confidence chance decoding interval). **A-B** results for ACC corresponding to Figure 3 **A-B** in main-text. **C-D** results for ACC corresponding to Figure 3 **E-F** in main-text.

**Figure S6.**
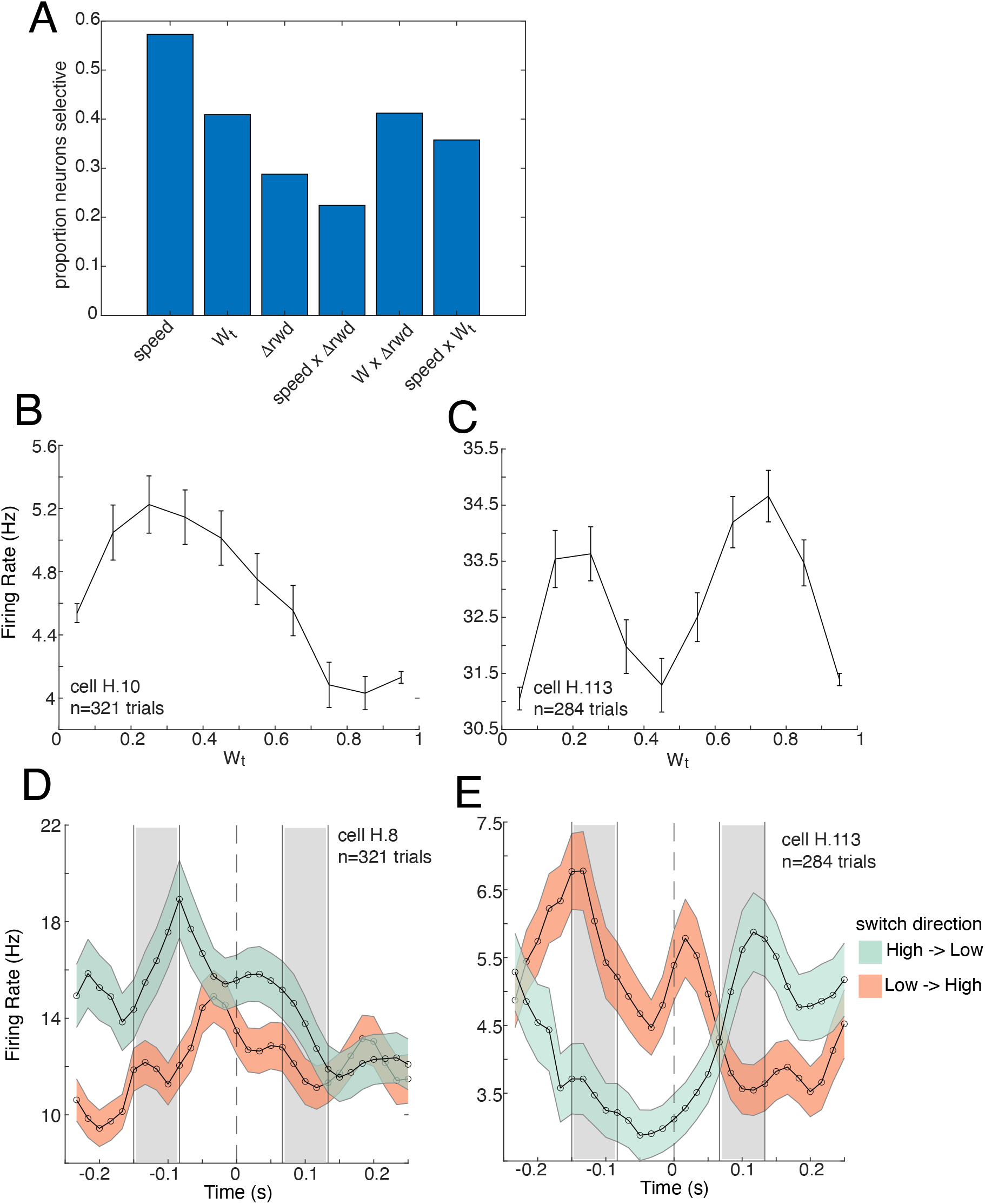
**A**. PMd selectivity results for linear regression model. **B-C** example PMd neuron tuning curves for *W*_*t*_, and **D-E** example neuron tuning over the change of mind window. In **D-E**, because the data are time-normalized and warped to the switch center (dashed line, W_*t*_=0.5), the gray regions denote the median switch-start and stop-time ± 1 std.

## REFERENCES

Acerbi, L., & Ma, W. J. (2017). Practical Bayesian optimization for model fitting with Bayesian adaptive direct search. Advances in neural information processing systems, 30.

Anderson, D. J., & Perona, P. (2014). Toward a science of computational ethology. Neuron, 84(1), 18–31.

Bishop, C. M. (2006). Pattern recognition and machine learning. Springer google schola, 2, 5–43.

Blanchard, T. C., Wolfe, L. S., Vlaev, I., Winston, J. S., & Hayden, B. Y. (2014). Biases in preferences for sequences of outcomes in monkeys. Cognition, 130(3), 289–299.

Blanchard, T. C., & Hayden, B. Y. (2014). Neurons in dorsal anterior cingulate cortex signal postdecisional variables in a foraging task. Journal of Neuroscience, 34(2), 646–655.

Brown, A. E., & De Bivort, B. (2018). Ethology as a physical science. Nature Physics, 14(7), 653–657.

Budišić, M., Mohr, R., & Mezić, I. (2012). Applied koopmanism. Chaos: An Interdisciplinary Journal of Nonlinear Science, 22(4).

Calhoun, A. J., Pillow, J. W., & Murthy, M. (2019). Unsupervised identification of the internal states that shape natural behavior. Nature neuroscience, 22(12), 2040–2049.

Cervera, R. L., Wang, M. Z., & Hayden, B. Y. (2020). Systems neuroscience of curiosity. Current Opinion in Behavioral Sciences, 35, 48–55.

Cisek, P. (2012). Making decisions through a distributed consensus. Current opinion in neurobiology, 22(6), 927–936.

Cisek, P., & Hayden, B. Y. (2022). Neuroscience needs evolution. Philosophical Transactions of the Royal Society B, 377(1844), 20200518.

Cooper, W. E., & Blumstein, D. T. (Eds.). (2015). Escaping from predators: an integrative view of escape decisions. Cambridge University Press.

Driscoll, L., Shenoy, K., & Sussillo, D. (2022). Flexible multitask computation in recurrent networks utilizes shared dynamical motifs. bioRxiv, 2022–08.

Dvijotham, K., & Todorov, E. (2012). A unifying framework for linearly solvable control. arXiv preprint arXiv:1202.3715.

Ebitz, R. B., & Hayden, B. Y. (2021). The population doctrine in cognitive neuroscience. Neuron, 109(19), 3055–3068.

Fieseler, C., Zimmer, M., & Kutz, J. N. (2020). Unsupervised learning of control signals and their encodings in Caenorhabditis elegans whole-brain recordings. Journal of the Royal Society Interface, 17(173), 20200459.

Fine, J. M., & Hayden, B. Y. (2022). The whole prefrontal cortex is premotor cortex. Philosophical Transactions of the Royal Society B, 377(1844), 20200524.

Fleming, S. M., Van Der Putten, E. J., & Daw, N. D. (2018). Neural mediators of changes of mind about perceptual decisions. Nature neuroscience, 21(4), 617–624.

Galgali, A. R., Sahani, M., & Mante, V. (2023). Residual dynamics resolves recurrent contributions to neural computation. Nature Neuroscience, 26(2), 326–338.

Hayden, B. Y. (2019). Why has evolution not selected for perfect self-control? Philosophical Transactions of the Royal Society B, 374(1766), 20180139.

Hayden, B. Y., Park, H. S., & Zimmermann, J. (2022). Automated pose estimation in primates. American journal of primatology, 84(10), e23348.

Hoeting, J. A., Madigan, D., Raftery, A. E., & Volinsky, C. T. (1999). Bayesian model averaging: a tutorial (with comments by M. Clyde, David Draper and EI George, and a rejoinder by the authors. Statistical science, 14(4), 382–417.

Khona, M., & Fiete, I. R. (2022). Attractor and integrator networks in the brain. Nature Reviews Neuroscience, 23(12), 744–766.

Kiani, R., Cueva, C. J., Reppas, J. B., & Newsome, W. T. (2014). Dynamics of neural population responses in prefrontal cortex indicate changes of mind on single trials. Current Biology, 24(13), 1542–1547.

Kobak, D., Brendel, W., Constantinidis, C., Feierstein, C. E., Kepecs, A., Mainen, Z. F., … & Machens, C. K. (2016). Demixed principal component analysis of neural population data. elife, 5, e10989.

Kutz, J. N., Brunton, S. L., Brunton, B. W., & Proctor, J. L. (2016). Dynamic mode decomposition: data-driven modeling of complex systems. Society for Industrial and Applied Mathematics.

Kwon, M., Daptardar, S., Schrater, P. R., & Pitkow, X. (2020). Inverse rational control with partially observable continuous nonlinear dynamics. Advances in neural information processing systems, 33, 7898–7909.

Lieder, F., & Griffiths, T. L. (2017). Strategy selection as rational metareasoning. Psychological review, 124(6), 762.

Linderman, S., Johnson, M., Miller, A., Adams, R., Blei, D., & Paninski, L. (2017, April). Bayesian learning and inference in recurrent switching linear dynamical systems. In Artificial Intelligence and Statistics (pp. 914–922). PMLR.

Markowitz, J. E., Gillis, W. F., Beron, C. C., Neufeld, S. Q., Robertson, K., Bhagat, N. D., … & Datta, S. R. (2018). The striatum organizes 3D behavior via moment-to-moment action selection. Cell, 174(1), 44–58.

Marques, J. C., Li, M., Schaak, D., Robson, D. N., & Li, J. M. (2020). Internal state dynamics shape brainwide activity and foraging behaviour. Nature, 577(7789), 239–243.

Murphy, K. P. (2022). Probabilistic machine learning: an introduction. MIT press.

Nair, A., Karigo, T., Yang, B., Ganguli, S., Schnitzer, M. J., Linderman, S. W., … & Kennedy, A. (2023). An approximate line attractor in the hypothalamus encodes an aggressive state. Cell, 186(1), 178–193.

Ostrow, M., Eisen, A., Kozachkov, L., & Fiete, I. (2024). Beyond geometry: Comparing the temporal structure of computation in neural circuits with dynamical similarity analysis. Advances in Neural Information Processing Systems, 36.

Pearson, J. M., Watson, K. K., & Platt, M. L. (2014). Decision making: the neuroethological turn. Neuron, 82(5), 950–965.

Pereira, T. D., Shaevitz, J. W., & Murthy, M. (2020). Quantifying behavior to understand the brain. Nature neuroscience, 23(12), 1537–1549.

Proctor, J. L., Brunton, S. L., & Kutz, J. N. (2016). Dynamic mode decomposition with control. SIAM Journal on Applied Dynamical Systems, 15(1), 142–161.

Rigotti, M., Barak, O., Warden, M. R., Wang, X. J., Daw, N. D., Miller, E. K., & Fusi, S. (2013). The importance of mixed selectivity in complex cognitive tasks. Nature, 497(7451), 585–590.

Sarafyazd, M., & Jazayeri, M. (2019). Hierarchical reasoning by neural circuits in the frontal cortex. Science, 364(6441), eaav8911.

Thura, D., & Cisek, P. (2016). Modulation of premotor and primary motor cortical activity during volitional adjustments of speed-accuracy trade-offs. Journal of Neuroscience, 36(3), 938–956.

Todorov, E. (2009). Compositionality of optimal control laws. Advances in neural information processing systems, 22.

Voloh, B., Maisson, D. J. N., Cervera, R. L., Conover, I., Zambre, M., Hayden, B., & Zimmermann, J. (2023). Hierarchical action encoding in prefrontal cortex of freely moving macaques. Cell reports, 42(9)

Wang, M. Z., & Hayden, B. Y. (2021). Latent learning, cognitive maps, and curiosity. Current Opinion in Behavioral Sciences, 38, 1–7.

Widge, A. S., Heilbronner, S. R., & Hayden, B. Y. (2019). Prefrontal cortex and cognitive control: new insights from human electrophysiology. F1000Research, 8.

Yeung, N., & Summerfield, C. (2012). Metacognition in human decision-making: confidence and error monitoring. Philosophical Transactions of the Royal Society B: Biological Sciences, 367(1594), 1310–1321.

Yoo, S. B. M., Hayden, B. Y., & Pearson, J. M. (2021). Continuous decisions. Philosophical Transactions of the Royal Society B, 376(1819), 20190664.

Yoo, S. B. M., Tu, J. C., Piantadosi, S. T., & Hayden, B. Y. (2020). The neural basis of predictive pursuit. Nature neuroscience, 23(2), 252–259.

Yoo, S. B. M., Tu, J. C., & Hayden, B. Y. (2021). Multicentric tracking of multiple agents by anterior cingulate cortex during pursuit and evasion. Nature communications, 12(1), 1985.

Yoo, S. B. M., & Hayden, B. Y. (2020). The transition from evaluation to selection involves neural subspace reorganization in core reward regions. Neuron, 105(4), 712–724.

